# RADseq data reveal a lack of admixture in a mouse lemur contact zone contrary to previous microsatellite results

**DOI:** 10.1101/2021.08.12.455854

**Authors:** Jelmer Poelstra, B. Karina Montero, Jan Lüdemann, Ziheng Yang, S. Jacques Rakotondranary, Paul Hohenlohe, Nadine Stetter, Jörg U. Ganzhorn, Anne D. Yoder

**Affiliations:** Department of Biology, Duke University, Durham, NC 27708, USA; Molecular and Cellular Imaging Center, Ohio State University, Wooster, OH 44691, USA; Institute of Zoology, Dept. Animal Ecology and Conservation, Universität Hamburg, 20146 Hamburg, Germany; Department of Genetics, Evolution and Environment, University College London, London, UK; Département Biologie Animale, Faculté des Sciences, P.O. Box 906, Université d’Antananarivo, Antananarivo, 101, Madagascar; Institute for Bioinformatics and Evolutionary Studies, Department of Biological Sciences, University of Idaho, Moscow, ID 83844, USA; Bernhard Nocht Institute for Tropical Medicine, 20359 Hamburg

**Author notes:** Co-senior authors (Jörg U. Ganzhorn, Anne D. Yoder). <u>Author contributions</u>: - Conception and design of study: - Data collection: JR, KM, NS, and JG collected samples in the field. PH and JP generated sequencing data. - Data analysis and interpretation: JP, ZY, JL, and KM performed all analyses. - Drafting and revising manuscript: JP, KM, JG, and ADY drafted the manuscript. All co-authors revised and agreed on the last version of the manuscript.

## Abstract

Despite being one of the most fundamental biological processes, the process of speciation remains poorly understood in many groups of organisms. Mouse lemurs are a species-rich genus of small primates endemic to Madagascar, whose diversity has only recently been uncovered using genetic data and is primarily found among morphologically cryptic, allopatric populations. To assess to what extent described species represent reproductively isolated entities, studies are needed in areas where mouse lemur taxa come into contact. Hybridization has previously been reported in a contact zone between two closely related mouse lemur species (*Microcebus murinus* and *M. griseorufus*) based on microsatellite data. Here, we revisit this system using RADseq data for populations in, near, and far from the contact zone, including many of the individuals that had previously been identified as hybrids. Surprisingly, we find no evidence for admixed nuclear ancestry in any of the individuals. Re-analyses of microsatellite data and simulations suggest that previously inferred hybrids were false positives and that the program `NewHybrids` can be particularly sensitive to erroneously inferring hybrid ancestry. Using coalescent-bases analyses, we also show an overall lack of recent gene flow between the two species, and low levels of ancestral gene flow. Combined with evidence for local syntopic occurrence, these data indicate that *M. murinus* and *M. griseorufus* are reproductively isolated. Finally, we estimate that they diverged less than a million years ago, suggesting that completion of speciation is relatively rapid in mouse lemurs. Future work should focus on the underpinnings of reproductive isolation in this cryptic primate radiation, which are mostly unknown. Our study also provides a cautionary tale for the inference of hybridization with microsatellite data.

## Introduction

Secondary contact zones, in which previously isolated populations meet, provide outstanding possibilities to investigate the mechanisms by which biodiversity accumulates. If contact zones form when reproductive isolation is incomplete, several outcomes are possible: the divergent populations can merge back together (e.g. Kearns et al. 2018), hybrid zones can form (Hewitt 2000, Hewitt 2001), and/or reinforcement of reproductive barriers can take place (e.g. Hoskin et al. 2005; Hopkins and Rausher 2012). The study of these different outcomes has contributed significantly our current understanding of speciation. For instance, hybrid zones offer opportunities to reveal the underlying basis of sources of reproductive isolation such as divergent phenotypes and genetic incompatibilities (Payseur 2010; Knief et al. 2019; Powell et al. 2020).

If secondary contact happens when reproductive isolation is complete, another set of outcomes is possible. One species may out-compete the other, preventing local overlap (e.g. Gurnell et al. 2004; Perry et al. 2007). Similarly, broad-scale mutual competitive exclusion and distinct habitat preferences may lead to adjacent yet largely non-overlapping distributions (Case et al. 2004; Wisz et al. 2012). Alpha biodiversity will only increase if species are able to co-occur locally, for instance by means of small-scale habitat heterogeneity or habitat partitioning among species, possibly after character displacement (e.g. Morris 1996; Arlettaz 2001; Estevo et al. 2017; Wuesthoff et al. 2021).

Mouse lemurs (genus *Microcebus*) provide an excellent organismal system for investigating these potential consequences of secondary contact. They are the world's smallest primates and are endemic to and widespread throughout Madagascar comprising as many as 25 named species, one of which was described as recently as 2020 (Schüßler et al. 2020). Additionally, there are several unnamed, hypothesized species (e.g. Louis et al. 2006). Species descriptions have relied heavily on genetic data because most species are morphologically highly cryptic and occur allopatrically, especially closely related ones (see Setash et al. 2017). Moreover, for many species descriptions, only mtDNA sequences have been analyzed and samples originated from a single or very few locations (Tattersall 2007 and references therein).

The combination of limited genetic and geographical sampling, little morphological differentiation, and allopatric occurrence has lead several authors to argue that the genus is likely to have been oversplit, possibly substantially so (Tattersall 2007; Markolf et al. 2011). One concern is that mtDNA divergence may not accurately reflect species divergence given that mtDNA represents only a single non-recombining, maternally inherited locus. This issue can be further exacerbated by limited geographic sampling which can cause clinal variation to be misinterpreted as the occurrence of multiple distinct clusters. Another concern is that lineages, even when they are indeed genetically divergent, may be more appropriately considered intraspecific variation (see also Coates et al. 2018). While examination with numerous nuclear loci and dense geographic sampling awaits for many mouse lemur species, so far, two studies have used RADseq data finding that genomic divergence largely (though not fully) corresponded to nominal species and mtDNA lineages (Yoder et al. 2016; Poelstra et al. 2020). The second concern, that lineages may be best described as distinct populations or subspecies, is harder to address. The fact that genomic data can be leveraged to obtain more accurate estimates of divergence times and rates of gene flow, and thus inform modern species delimitation analyses (e.g. Poelstra et al. 2020; Dincă et al. 2019; Hundsdoerfer et al. 2019), does offer the promise of a more nuanced assessment of taxonomic boundaries.

Even so, there are distinct limits to this approach when lineages occur allopatrically given that the key measure of speciation – whether and to what extent reproductively isolation (RI) exists between divergent lineages – cannot be directly assessed in the absence of experimental approaches that are time-consuming and not feasible for many non-model organisms. Thus, studies of divergent lineages in secondary contact are needed to gain insight into types and levels of divergence that do or do not produce RI. From a practical species delimitation perspective, this will also allow for the comparative examination of divergence for allopatric lineages.

To date, seven different pairs of mouse lemur species have been shown to co-occur locally at various localities throughout Madagascar. One widespread species, *M. murinus*, is involved in five of these cases. *M. murinus* co-occurs with its sister species *M. griseorufus* in southern Madagascar and from south to north in western Madagascar with *M. berthae*, *M. myoxinus*, *M. ravelobensis*, and *M. bongolavensis*, respectively (Radespiel 2016; Sgarlata et al. 2019; Wuesthoff et al. 2021). In northeastern Madagascar, two other species pairs occur in local sympatry: *M. mittermeieri* and *M. macarthurii* (Radespiel et al. 2008; Poelstra et al. 2020) as well as *M. lehilahytsara* and *M. jonahi* (Poelstra et al. 2020; Schüßler et al. 2020).

In all but one of these seven cases of sympatry, no hybridization has been detected Sources of reproductive isolation among sympatric mouse lemurs are poorly known, but factors that may contribute to prezygotic isolation via differential mate choice may include divergence in acoustic (Braune et al. 2008; Hasiniaina et al. 2020) and olfactory signaling (Kollikowski et al. 2019; Hunnicutt et al. 2020). Additionally, opportunities for reproductive interaction may be reduced by ecological divergence manifesting, for example, in differential timing of the highly seasonal and temporally constrained reproductive season seen in mouse lemurs (Schmelting 2000; Evasoa et al. 2018; Schüßler et al. 2020).

Hybridization has only been detected between *M. murinus* and *M. griseorufus* (Gligor et al. 2009; Hapke et al. 2011), which is also unique among the seven cases of sympatry in consisting of a pair of sister lineages. It should be noted that populations of *M. murinus* studied by Gligor et al. (2009) have since been split as *M. manitatra* and *M. ganzhorni* by Hotaling et al. (2016), whereas the populations studied by Hapke et al. (2011) continue to be part of *M. murinus*. However, we here include *M. manitatra* and *M. ganzhorni* under the nomer “*M. murinus s.l.*” (Weisrock et al. 2010) pending further taxonomic revisions (see Methods for further details).

Gligor et al. (2009) studied an area where *M*. *murinus s.l.* and *M*. *griseorufus* come into geographic contact. They sequenced part of the mitochondrial HV1 locus and genotyped 9 microsatellite loci for a total of 162 mouse lemurs at three spiny forest sites with *M*. *griseorufus* (n=26), three littoral forest sites for *M*. *murinus* (n=98), and three sites in an ecotone between spiny and littoral forest with both species (n=38). Using the programs `STRUCTURE` (Pritchard et al. 2000) and `GeneClass` (Piry et al. 2004), they concluded that “most individuals within the transition zone” had mixed ancestry (no individual-level assignments were made). Hapke et al. (2011) studied a contact zone 40 km further north, where, instead of a gradual transition between habitat types, narrow strips of mesic gallery forest along rivers and streams directly border dry spiny forest in the surrounding areas. This study used the same set of microsatellite loci for a total of 159 mouse lemurs, with `STRUCTURE` and `NewHybrids` (Anderson and Thompson 2002) identifying a total of 18 admixed individuals, originating from all but one of the six sites examined (highest percentage of hybrids: 17.3% out of 75 at Mangatsiaka). Of these, 15 individuals showed signs of nuclear admixture (i.e., among microsatellites) whereas 3 had a mismatch between microsatellite and mitochondrial ancestry.

Based on the results of Gligor et al. (2009) and Hapke et al. (2011), the contact zones between these species seemed to provide an ideal opportunity for studying species separation based on species-specific microhabitat utilization, and its breakdown along ecotones or in disturbed areas where habitat patches become too small to allow for habitat-specific separation (Rakotondranary and Ganzhorn 2011; Rakotondranary et al. 2011). In a follow-up study (Sommer et al. 2014), hybrids showed a higher prevalence of intestinal parasites, and several MHC alleles were found to be shared between both species and their putative hybrids.

Here, we revisit the contact zone area studied by Hapke et al. (2011) using RADseq data. We have included many of the individuals that were inferred to be hybrids by that study in addition to samples from nearby and distant allopatric populations. We examined individual-level admixture in the contact zone and used coalescent modeling to ask whether there is evidence for ongoing and or ancestral gene flow between the species. To our surprise, we found no evidence for admixed individuals in the contact zone – including among the individuals previously identified as hybrids – and have also inferred a lack of ongoing gene flow between the two species more generally.

## Methods

### Sampling

Hapke et al. (2011) and follow-up work in Lüdemann (2018) detected hybridization between *M. murinus* (hereafter referred to as *murinus*) and *M. griseorufus* (hereafter referred to as *griseorufus*) using 9 microsatellites and a fragment of the HV1 mitochondrial locus from individuals in the Andohahela area in southeastern Madagascar. We made use of a selection of their samples, for which trapping and sample collection procedures are described in Gligor et al. (2009), Hapke et al. (2011), and Lüdemann (2018). We augmented this dataset with 13 *griseorufus* and 20 *murinus s.l.* (see below) samples from distant, allopatric sites, and with 3 *M. rufus* samples that were used as an outgroup. Ear clips from wild-caught and released mouse lemurs were collected between 2006 and 2017 (*Table S1*, *Table S2*).

At two of the six sites examined by Hapke et al. (2011), they detected unadmixed individuals of both parental species as well as individuals with admixed ancestry (individuals inferred to be admixed by Hapke et al. (2011) and Lüdemann (2018) are hereafter referred to as “putative hybrids”). Given this community composition, we refer to these two contact zone sites, Mangatsiaka and Tsimelahy, which are ~6.5 kilometers apart, as “sympatric” sites. From these two sites, we selected 78 samples for the present study (*Table S1*). Among the 49 samples from Mangatsiaka that we sequenced, Hapke et al. (2011) and Lüdemann (2018) classified 21 as *murinus* based on microsatellites as well as mtDNA, 13 as *griseorufus* based on microsatellites as well as mtDNA, 3 as *griseorufus* based on microsatellites but as *murinus* based on mtDNA (i.e. these individuals had a mitonuclear ancestry mismatch), and 14 as admixed based on the microsatellites (i.e. putative hybrids, of which 7 had a *griseorufus* mtDNA haplotype, and 7 had a *murinus* mtDNA haplotype). Among the 29 individuals from Tsimelahy, Hapke et al. (2011) classified 15 as pure *murinus*, 15 as pure *griseorufus*, and 1 as admixed based on microsatellites (this individual had a *griseorufus* mtDNA haplotype). Thus, in total, we sequenced 15 individuals for which Hapke et al. (2011) or Lüdemann (2018) had detected nuclear admixture, and an additional 3 with a mitonuclear ancestry mismatch.

We additionally selected samples from nearby sites at which Hapke et al. (2011) had exclusively (or nearly so) detected unadmixed individuals of one of the two species: 8 *griseorufus* from Hazofotsy and 8 *murinus* from Ambatoabo (*Table S1*). We refer to these contact zone sites as “parapatric” sites. Hazofotsy is 14.5 kilometers from Mangatsiaka, whereas Ambatoabo is 14 kilometers from Tsimelahy (*Fig. 1* - inset). In total, we sequenced 94 samples from the contact zone area (sympatric and parapatric sites) in the Andohahela area.

**Fig. 1:**
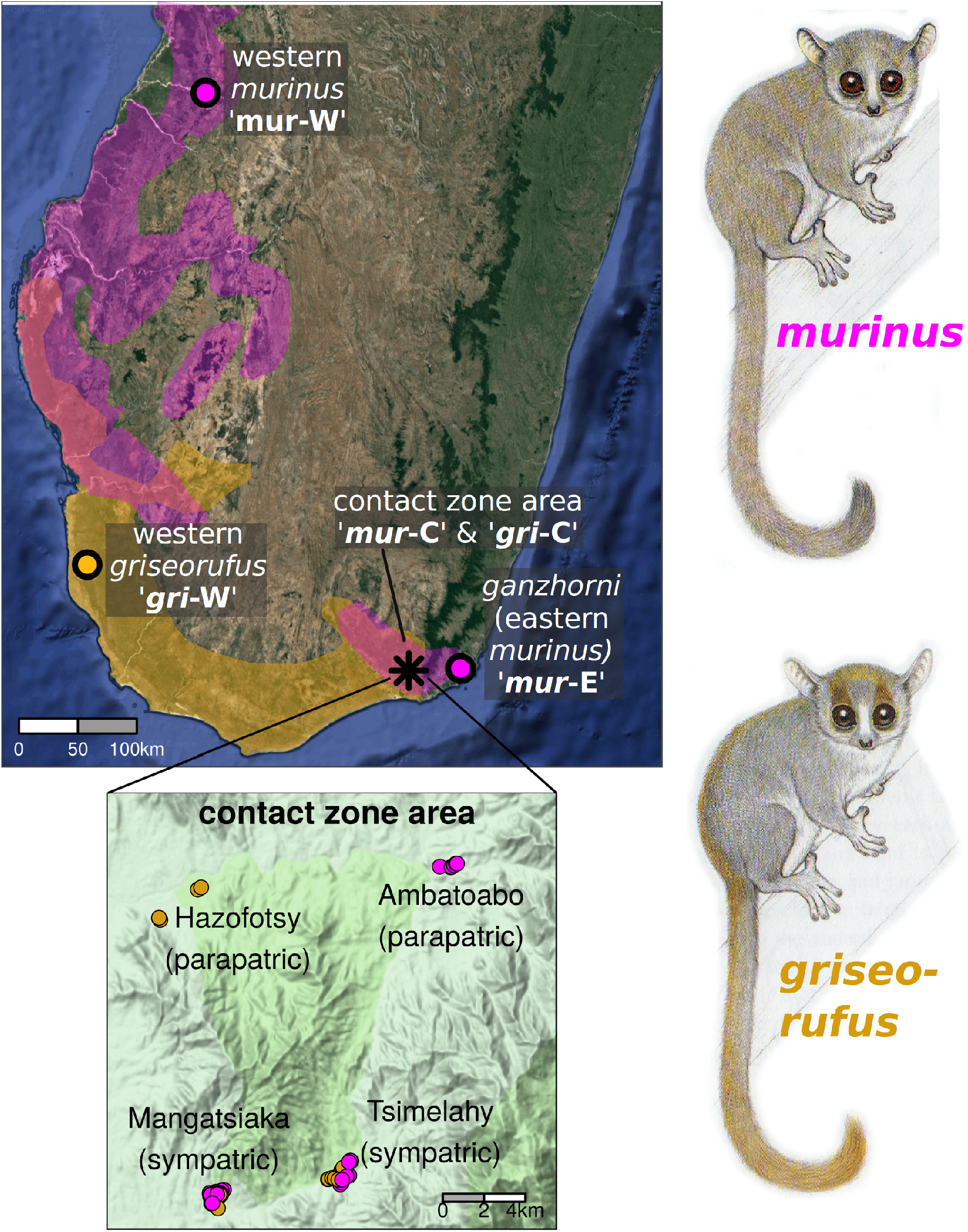
Distributions and sampling sites of *murinus* and *griseorufus* in southern Madagascar. The distribution of *murinus* is shown in purple and that of *griseorufus* in gold. A population in southeastern Madagascar was recently split from *murinus* as *M. ganzhorni*, but is here included within *murinus* s.l‥ A large gap across the central part of southern Madagascar divided *murinus* populations, and sampling areas *mur*-C and *mur*-E are together referred to as “southeastern *murinus* populations”. Note that the range of *M. murinus* extends to the north of the area shown in the map, whereas the entire distribution of *M. griseorufus* is shown. **Inset**: Overview of sampling in the contact zone area, showing two parapatric (Hazofotsy with *griseorufus* and Ambatoaba with *murinus*) and two sympatric (Mangatsiaka and Tsimelahy) sites.

Finally, “allopatric” samples, taken well away from the contact zone, were represented by 14 *griseorufus* from several sites in southwestern Madagascar, 8 *murinus* from several sites in western Madagascar, and 11 *M. ganzhorni,* a species that was recently split from *murinus* (Hotaling et al. 2016), from Mandena in far southeastern Madagascar (*Table S2*, *Fig. 1*). Below, we show that *M. ganzhorni* diverged from the *murinus* populations from the Andohahela area very recently, while a much deeper split occurs between other populations from southeastern Madagascar and those from Madagascar that all continue to be classified as *murinus.* Therefore, as mentioned above, we here include *M. ganzhorni* (and *M. manitatra*, which was not included in this study) under the nomer “*M. murinus s.l.*”. As an outgroup, we used *M. rufus* (three samples, *Table S2*).

We used the following geographically defined population groupings for analyses where individuals are assigned to predefined groups (*Fig. 1*): western *griseorufus* (abbreviated “*gri*-W”), central/contact zone area *griseorufus* (abbreviated “*gri*-C*”)*, western *murinus* (abbreviated “*mur*-W”), central/contact zone area *murinus* (abbreviated “*mur*-C*”*), and eastern *murinus s.l.* (abbreviated “*mur*-E”; this population corresponds to *M. ganzhorni* sensu Hotaling et al. (2016), see details above). The *mur*-C and *mur*-E populations are geographically and phylogenetically close and are sometimes together referred to as “southeastern *murinus* populations”.

### Sequencing

We prepared Restriction-site Associated DNA (RAD) sequencing libraries using 50 ng of genomic DNA from each sample following the protocol of Ali et al. (2016). Briefly, samples were digested with SbfI (New England Biolabs), followed by ligation with custom biotinylated adapters containing 8 bp barcodes unique to each sample. We pooled 48 samples in a single library, with a technical replicate for four of these samples, and sheared DNA to an average fragment size of 400 bp using a Covaris M220. RAD fragments were enriched with a streptavidin bead pull-down and prepared as a sequencing library using a NEBNext Ultra DNA Library Prep Kit (New England Biolabs). Final libraries were sequenced using paired-end 150 bp sequencing on an Illumina HiSeq 4000 at Duke University's Center for Genomic sand Computational Biology sequencing facility.

### RADseq bioinformatics and genotyping

When using the Ali et al. (2016) protocol, half of the barcodes end up in the reverse (R2) reads. Therefore, raw reads in FASTQ files were first “flipped” using a custom Perl script, and were next demultiplexed and deduplicated in `Stacks` v2.0b (Rochette et al. 2019) using the “`process_radtags`” and “`clone_filter`” commands, respectively. Reads were then quality filtered using `Trimmomatic` (Bolger et al. 2014) with the following parameters: Leading: 3, Trailing: 3, Slidingwindow: 4:15, Minlen: 60. Reads were aligned to the *M. murinus* reference genome (“Mmurinus 3.0”, https://www.ncbi.nlm.nih.gov/genome/777?genome_assembly_id=308207, Larsen et al. 2017) with BWA MEM v0.7.15 (Li 2013). From the resulting BAM files, reads that were properly paired and had a minimum mapping quality of 30 were retained using “`samtools view`” (“-f 0×2” and “-q 30” arguments, respectively), and filtered BAM files were sorted using “`samtools sort`”, all from the `SAMtools` library (v1.6, Li et al. 2009).

We performed genotype calling with `GATK` v4.0.7.0 (DePristo et al. 2011), and we filtered SNPs and individuals largely according to the "FS6" filter of O‘Leary et al. (2018) (see *Supplementary Materials* for details). Unless otherwise noted, downstream analyses used sets of SNPs that resulted from this filtering procedure for all analyses except the coalescent-based modeling. The filtering procedure, which includes several consecutive rounds of removing the individuals and SNPs with the highest amounts of missing data, was performed separately for the set of all 135 sequenced individuals (including the 3 outgroup individuals; the resulting VCF was used for phylogenetic inference, admixture statistics, and served as the basic for generating full-sequence loci for coalescent-based modeling) and for the set of 94 individuals from the contact zone area (the resulting VCF was used for clustering analyses).

For the set of individuals from the contact zone area, we additionally produced two datasets using more lenient filtering procedures, to be able to examine admixture using more individuals and SNPs: (1) a dataset produced by omitting the last round of removal of SNPs and individuals based on missing data; (2) a dataset produced using the FS6 filter without the individual-filtering steps that retained two additional putative hybrids and two individuals with mitonuclear discordance.

We computed the following quality control statistics for each sample and then compared these between samples that had previously been identified as *murinus*, as *griseorufus*, or as hybrid: number of filtered FASTQ reads, depth of coverage in BAM files, mean mapping quality, percentage of reads that were mapped, percentage of reads that were properly paired, depth of coverage, and the percentage of missing data in VCF files.

Based on `GATK`-called genotypes, we also produced full-sequence FASTA files for each RAD locus (see *Supplementary Materials* for details).

### Detection of hybrids using clustering approaches

For the detection of admixed individuals, we used complementary model-free and model-based approaches. First, we used Principal Component Analysis (PCA) as implemented in the `SNPRelate R` package v1.17.2 (Zheng et al. 2012), using the `snpgdsPCA()` function (after conversion from the VCF file with the `snpgdsVCF2GDS()` and `snpgdsOpen()` functions). Second, we used the program `ADMIXTURE` v1.3.0 (Alexander et al. 2009) to detect clusters and assign individual-level ancestry proportions from each cluster. Third, we used the program `NewHybrids` v1.1 (Anderson and Thompson 2002), which identified the majority of admixed individuals in Hapke et al. (2011) and Lüdemann (2018). `NewHybrids` was used to estimate, for each sample, the posterior probability of it belonging to each of six predefined categories: *griseorufus*, *murinus*, F1 hybrid (*griseorufus* x *murinus*), F2 hybrid (F1 x F1), *griseorufus* backcross (F1 x *griseorufus*) and *murinus* backcross (F1 x *murinus*). 500,000 iterations were used as burn-in, with another 1,500,000 iterations after that, using Jaffereys-like priors. A run was considered successful if it passed a test for convergence implemented in the `hybriddetective R` package (Wringe et al. 2017).

These analyses were first performed with datasets produced by passing individuals only from the contact zone area (i.e., the sympatric and parapatric sites), through the three filtering procedures described above. In addition, we ran these analyses for a dataset produced by passing *all* individuals (i.e., including individuals from allopatric populations) through the standard genotyping filter.

### Reanalysis of microsatellite data

We reanalyzed the Hapke et al. (2011) and Lüdemann (2018) microsatellite data using only the samples included in this study. Like in Hapke et al. (2011), we used the Bayesian classification methods `STRUCTURE` v. 2.3.4 (Pritchard et al. 2000; see the *Supplementary Materials* for details) and `NewHybrids` v. 1.1 to detect hybrids. For `STRUCTURE`, 20 runs using K=2 were used to calculate the average membership coefficients by creating an optimal alignment using the full-search algorithm implemented in `CLUMPP` v. 1.1.2 (Jakobsson and Rosenberg 2007). To keep the results directly comparable, we used the same threshold for the detection of hybrids as Hapke et al. (2011): a sample was considered a hybrid when the posterior probability for assignment to the species of their mitochondrial haplotype was ≤ 0.9 for `Structure` or ≤ 0.5 in NewHybrids, and part of a specific hybrid category when the corresponding probability was > 0.5.

### Comparison of microsatellites and SNPs using simulations

Using simulations, we compared the performance of microsatellites and SNPs for detecting hybrids. The `hybriddetective` R package (Wringe et al. 2017) was used to generate multi-generational hybrids from both the microsatellite and SNP data. First, unadmixed *murinus* and *griseorufus* individuals were created by randomly drawing two alleles per locus from the allopatric reference populations, without replacement. For subsequent F1 samples, one allele per locus was drawn from an unadmixed individual of each species. This procedure, drawing from the appropriate population, was continued for F2 and backcross individuals. In total, 60 simulated individuals were created: 20 each of unadmixed *griseorufus* and *murinus*, and 5 each of F1, F2, F1 x unadmixed *griseorufus*, and F1 x unadmixed *griseorufus*. Ancestry assignment was compared between microsatellites and SNPs by running `STRUCTURE` and `NewHybrids`, as described above, on the simulated genotypes.

### Phylogenetic inference

To enable subsequent tests of gene flow and demographic modeling, we determined relationships among all *murinus s.l.* and *griseorufus* individuals sampled by our study, using three *M. rufus* individuals as an outgroup. These analyses also provided a first-pass exploration of patterns of gene flow.

First, we used the NeighborNet method implemented in `Splitstree` v. 4.14.4 (Huson and Bryant 2006). This method visually displays phylogenetic conflict in an unrooted tree and thus shows phylogenetic relationships while also allowing for the detection of potentially admixed populations and individuals.

Second, we used `Treemix` v1.13 (Pickrell and Pritchard 2012) to estimate relationships among predefined populations (*gri*-W, *gri*-C, *mur*-W, *mur*-C, and *mur*-E) both with and without admixture events among populations, which are inferred based on a user-defined number of admixture events. We used a number of admixture events *m* ranging from 0 to 10, and 100 bootstraps. We performed likelihood-ratio tests to determine the most likely number of migration events, comparing each graph to one with one fewer migration event, and took the first non-significant comparison as the most likely number of migration events.

### Formal admixture statistics

The D-statistic and related formal statistics for admixture use phylogenetic invariants to infer post-divergence gene flow between non-sister populations. We used the `qpDstat` program of `admixtools` v4.1 (Patterson et al. 2012) to compute four-taxon D-statistics, which test for gene flow between P3 and either P1 or P2, given the tree topology (((P1, P2), P3), P4).

We used all possible configurations in which gene flow between *murinus* and *griseorufus* could be detected. First, we used the five main populations (*gri*-W, *gri*-C, *mur*-W, *mur*-C, *mur*-E). In order to test for admixture limited to the specific sites where contact between the two species currently occurs, we next divided the contact zone area populations (*gri*-C, *mur*-C) into two groups each: sympatric (Mangatsiaka and Tsimelahy) and parapatric sites (Ambatoabo and Hazofotsy, see *Fig. 1* - inset). For all tests, *M. rufus* was used as P4 (the outgroup). Significance of D-values was determined using the default Z-value reported by `qpDstat`, which uses weighted block jackknifing. This approach to determining significance is conservative for RADseq data given that linkage disequilibrium is, on average, expected to be lower between a pair of RADseq SNPs than between a pair of SNPs derived from whole-genome sequencing (Patterson et al. 2012; Kim et al. 2018).

Admixture proportions can be estimated using f_4_-ratio tests for tree topologies using five populations wherein P_x_ is potentially admixed between P2 and P3, with P2 sister to P1, and O as an outgroup to the other four populations. Using this framework, we first tested whether and to what extent contact zone area populations of either species (*mur*-C and *gri*-C) are admixed with one more populations of the other species, with follow-up tests using *mur*-E as the potentially admixed population P_x_.

### Demographic Modeling

`First, G-PhoCS` v1.3 (Gronau et al. 2011), a coalescent-based approach that utilizes Markov Chain Monte Carlo (MCMC) was used to jointly infer population sizes, divergence times and migration rates for the three *murinus* populations (*mur*-W, *mur*-C, and *mur*-SE) and the two *griseorufus* populations (*gri*-W and *gri*-SE). Because it was not computationally feasible to run `G-PhoCS` for the entire dataset, we selected, for each population, 3 individuals that had high coverage and low amounts of missing data, while ensuring that mean coverage and missing data amounts were approximately equal across populations. Because `G-PhoCS` does not infer phylogenetic relationships among populations, the species tree recovered from phylogenetic analyses (above) was fixed for parameter sampling. As input, we created full-sequence FASTA files based on the `GATK` genotypes (See *Supplementary Materials* for details).

Gene flow is modelled in `G-PhoCS` using one or more discrete unidirectional migration bands between a pair of extant or ancestral lineages that overlap in time. Since each migration band adds a parameter to the model, it is often not feasible to include all possible migration bands. Here, we modelled reciprocal migration bands between *gri*-C and *mur*-C and between ancestral *griseorufus* and *murinus* lineages, as we were interested in the occurrence gene flow between *griseorufus* and *murinus* in the contact zone and in more ancient gene flow between the two species. Additionally, we ran a model with no migration bands to assess how this affected divergence time and population size estimates.

Second, we ran the multispecies-coalescent-with-introgression (MSCi) model in `BPP` v. 4.2 (Flouri et al. 2020) using the same set of full-sequence loci used for `G-PhoCS`. While `G-PhoCS` implements an isolation-with-migration model with continuous gene flow during potentially long periods, the MSCi model in `BPP` models discrete introgression events. For each introgression event, it estimates the introgression probability φ, which represents the proportion of loci inherited from one of the two parents of an introgression node. We conducted 4 replicate runs all of which assessed support for 6 introgression events during the same periods for which gene flow was modelled in `G-PhoCS`: between the extant *mur*-C and *gri*-C populations, between the *murinus* lineage ancestral to *mur*-C and *mur*-E and the co-temporal ancestral *griseorufus* lineage, and between ancestral *murinus* and *griseorufus* lineages prior to intraspecific divergence in both species.

### Conversion of demographic parameters

We converted the migration rate parameter *m* to the population migration rate (2Nm), which is the number of haploid genomes (i.e., twice the number of migrants) in the source population that arrive each generation by migration from the target population. The population migration rate is calculated using the value of θ for the target population [2Nm_s→t_ = *m*_s→t_ × (θ_t_/4)], and as such it does not depend on an estimate of the mutation rate.

Divergence times, population sizes and the proportion of migrants per generation (*m* x μ) were converted using empirical estimates of the mutation rate and generation time. To incorporate uncertainty in these estimates, we drew a random number from distributions for the mutation rate and generation time for each sampled MCMC generation. We used a mutation rate of 1.52 x 10^-8^, which is the pedigree-based mutation rate estimate for *M. murinus* from Campbell et al. (2021). For the generation time, we used a lognormal distribution with a mean of ln(3.5) and a standard deviation of ln(1.16) based on two available estimates for *Microcebus* (4.5 years from Zohdy et al. 2014 and 2.5 years from Radespiel et al. 2019).

## Results

### Genotyping and QC statistics

`GATK` genotyping followed by the standard (“FS6”) filtering procedure for all individuals resulted in a VCF file with 79 individuals and 56,255 SNPs. The equivalent VCF file with only samples from sympatric and parapatric sites in the contact zone area (Andahohela area, see *Fig. 1*) contained 69 individuals, 12 of which were putative hybrids, and 7,180 SNPs. The two less stringent filtering procedures (see Methods) for the contact zone set resulted in the retention of 78 individuals (13 putative hybrids) and 48,556 SNPs and 79 individuals (18 putative hybrids) and 1,360 SNPs, respectively. 16 individuals, among which 2 putative hybrids, did not survive the filtering steps for any of the final VCF files.

The full-sequence FASTA file produced for `G-PhoCS` analyses contained 12,952 loci with an average length of 475 bp.

QC statistics were overall highly similar between *murinus*, *griseorufus*, and putative hybrid samples from the contact zone area (*Fig. S1-S10*, *Table S3*). Statistics related to read mapping were slightly lower for *griseorufus* than for *murinus*, which is expected given that the reference genome is *murinus*: the percentage of mapped reads (means of 93.4% and 93.9%, respectively; *Fig. S4*), the mean mapping quality for unfiltered BAM files (means of 44.6 and 45.8, respectively; *Fig. S5*). For these statistics, putative hybrids were intermediate, which would be expected both if they were true hybrids and if they consisted of a mixture of individuals from either species. The percentage of properly paired reads differed very little between *griseorufus* (99.76%) and *murinus* (99.85%), though these distributions barely overlapped and putative hybrids separated in two clusters (*Fig. S6*).

A lower percentage of *griseorufus* samples passed the standard filtering procedure (“FS6”, 60.5% vs 83.7% for *murinus*, *Table S3*) but for samples passing these filtering steps, mean depth and the percentage of missing SNPs were similar between the two species (mean depth: 39.8x for *griseorufus* and 38.2x for *murinus*; mean percentage of missing SNPs 2.25% for *griseorufus* and 2.49% for *murinus*; *Fig. S9-S10*, *Table S4*). While putative hybrids had a slightly lower depth (34.7x) and higher missingness (2.92%) in the final VCF (*Fig. S9-S10*, *Table S4*), the absolute values are no cause of concern for subsequent analyses.

### No evidence for ongoing hybridization in the contact zone

`ADMIXTURE` identified K=2 as the optimal number of clusters among individuals from the contact zone area (*Fig. 2A* - top). All individuals, including the 12 putative hybrids that passed filtering, were entirely assigned to one of the two clusters (*Fig. 2A* - bottom), with no signs of admixture. Results were also plotted for K=3, for which a third cluster corresponded to differentiation between sympatric (Mangatsiaka, Tsimelahy) and parapatric (Hazofotsy) sites in *griseorufus* (*Fig. S11*).

**Fig. 2:**
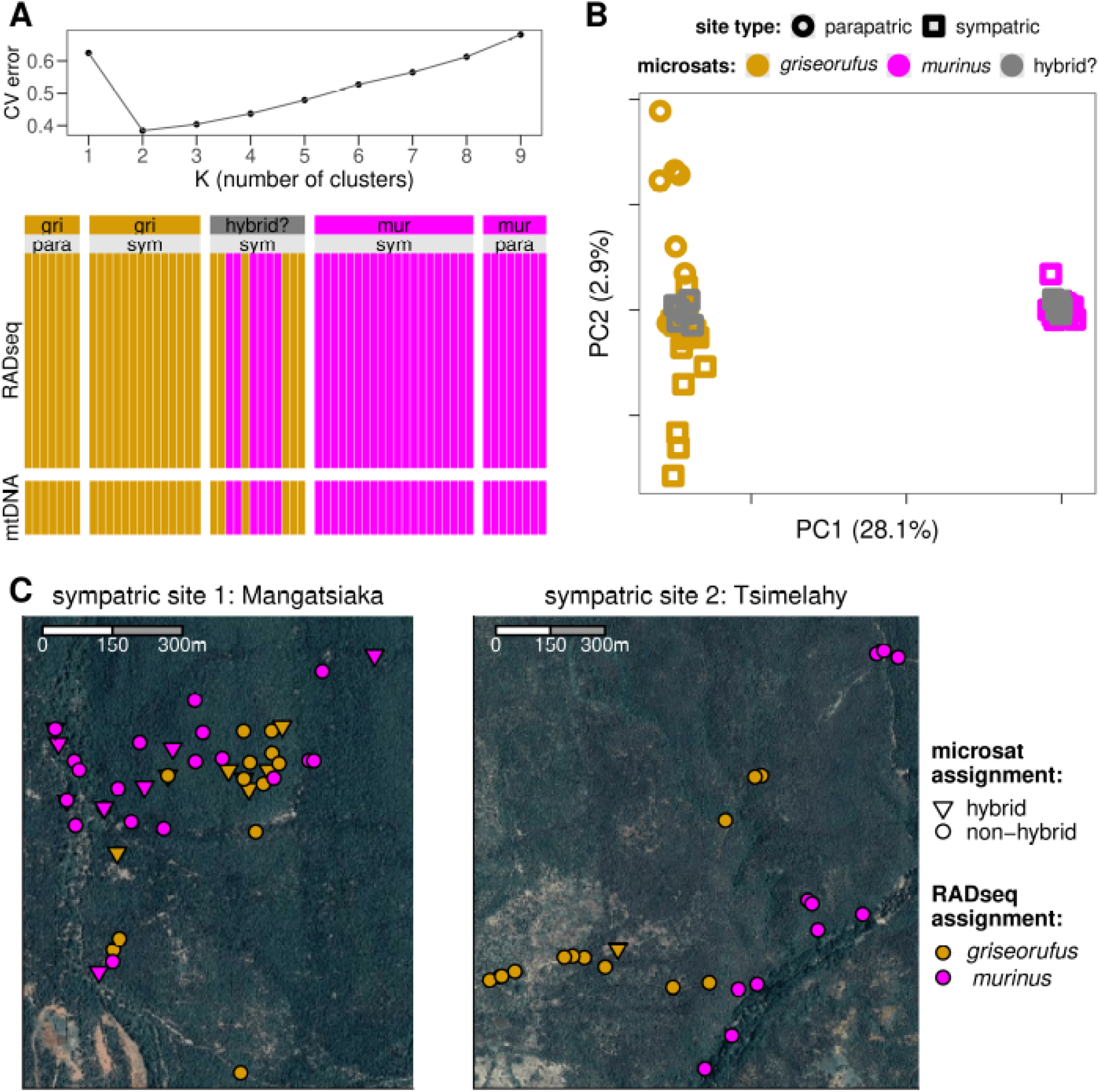
No evidence for hybridization in the contact zone. Nuclear RADseq data from the contact zone area was used for all analyses, including 12 individuals that had been identified as admixed in a previous microsatellite study (dark gray in panels A and B). **A)** ADMIXTURE results. Top: a cross-validation error plot identifies K=2 as the optimal number of clusters. Bottom: Ancestry components for each individual for K=2 reveal a lack of admixture: all individuals were inferred to have 100% ancestry from only a single species. Individuals were previously characterized using mtDNA (bottom bars) and microsatellites (labels at top). **B)** A PCA analysis reveals two clusters that are well-separated along PC1, corresponding to *griseorufus* and *murinus*, with no individuals that are intermediate along this axis. **C)** Map showing spatial distribution of *murinus* and *griseorufus* individuals at the two contact sites.

Principal component analysis (PCA) with individuals from the contact zone revealed a wide separation between two groups along the first principal component axis (PC1), which explained around tenfold more of the variation compared to PC2. The separation along PC1 corresponded to differentiation between *griseorufus* and *murinus,* and importantly, all putative hybrids fell within one of those two groups, with none occupying an intermediate position (*Fig. 2B*). Similar to the `ADMIXTURE` results at K=3, PC2 mostly corresponded to differentiation between sympatric and parapatric sites in *griseorufus* (see also *Fig. S12* for a within-species PCA).

`NewHybrids` was run with and without assigning individuals from the parapatric populations to reference parental species, and in both cases, all individuals were assigned to one of the two parental species and none were assigned to one of the hybrid categories. Assignment to species matched perfectly with `ADMIXTURE` assignments and PCA results.

Datasets produced by less stringent filtering procedures included an additional 4 putative hybrids that did not pass all filtering steps but could still be assessed using a more limited number of SNPs (*Fig. S13*). `ADMIXTURE` and `NewHybrids` analyses of these datasets similarly showed no evidence for admixed individuals with the exception of mitonuclear discordance: for two of the individuals for which Lüdemann (2018) had detected *griseorufus* ancestry in nuclear DNA but *murinus* mtDNA haplotypes mitonuclear discordance, we could confirm that the nuclear DNA has pure *griseorufus* ancestry (*Fig. S13*). The third sample for which Lüdemann (2018) detected mitonuclear discordance did not pass filtering at all. No other cases of mitonuclear discordance were found (Fig. 2A, *Table S1*.)

### False positives in hybrid detection using microsatellites with NewHybrids

In a reanalysis of the Hapke et al. (2011) microsatellite data for only the individuals that were included in this study, 11 individuals identified as hybrids in Hapke et al. (2011) were no longer identified as such by either `NewHybrids` or `STRUCTURE`. Only a single sample was now identified as a hybrid by `NewHybrids`, but `STRUCTURE` did not support this inference (*Fig. 3A*, *Fig. S14*). As noted above, admixture was not detected for any individuals in the RADseq data, including those that had been identified as hybrids in the original microsatellite analyses.

**Fig. 3:**
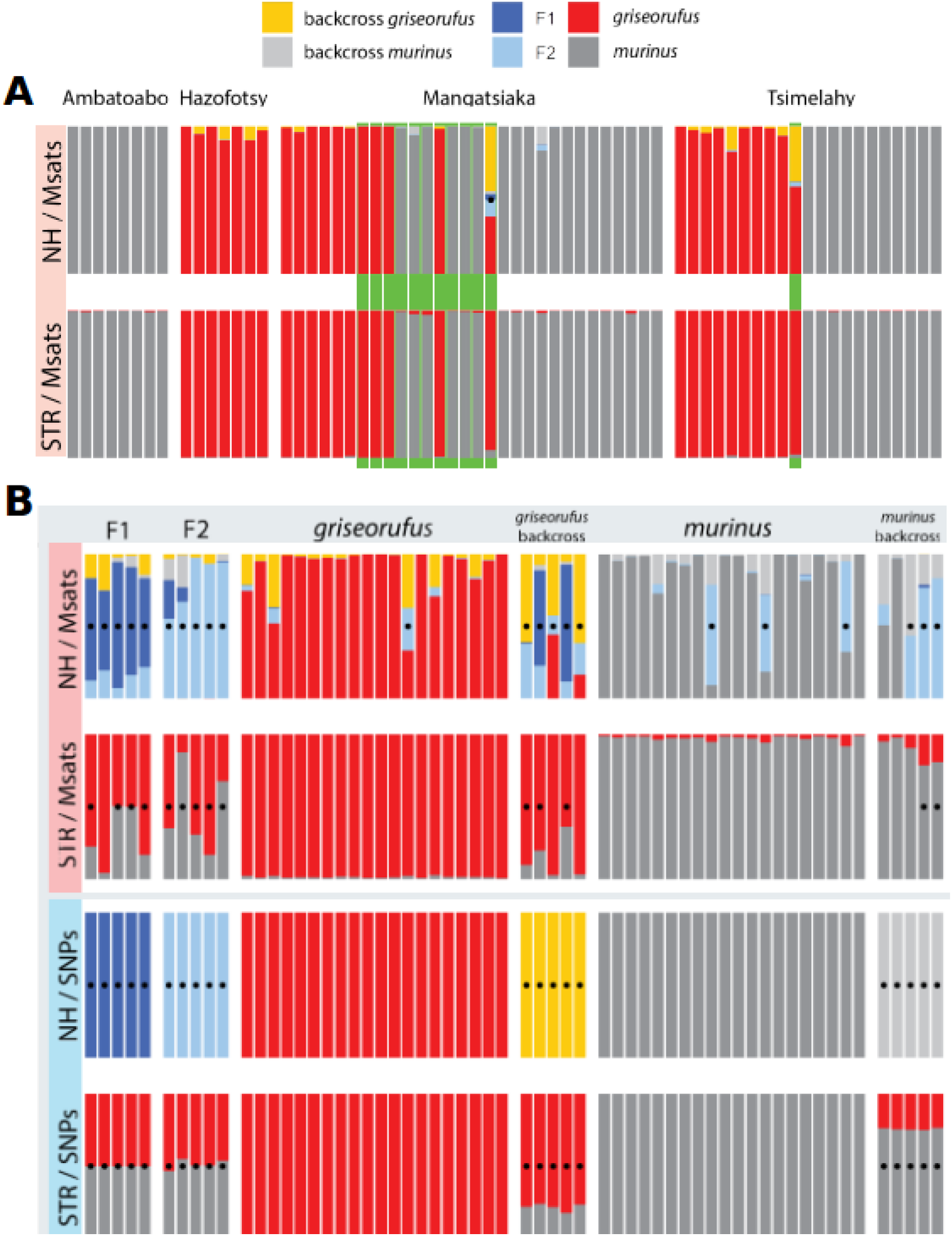
Re-analysis of microsatellite data and analysis of simulated individuals. **A)** Re-analysis of microsatellite data with `NewHybrids` (NH; top row) and `STRUCTURE` (STR; bottom row). Among the 12 individuals previously identified as hybrids (green background bars), `NewHybrids` now identifies only a single individual as a hybrid (black dot), with several further *griseorufus* individuals showing non-significant signs of admixed ancestry (yellow ancestry). **B)** Analysis of simulated individuals. Dots indicate detected hybrids. Using SNPs (bottom two rows), both `NewHybrids` and `STRUCTURE` correctly inferred ancestry for all individuals. Using microsatellites (top two rows), `NewHybrids` was prone to falsely inferring hybrids (4 out of 40 unadmixed individuals), and false negatives occurred both with `NewHybrids` (2 out of 20) and `STRUCTURE` (6 out of 20).

In analyses of simulated microsatellite data, `NewHybrids` inferred that 4 out of 40 unadmixed individuals were hybrids, whereas `STRUCTURE` found no false positives. False negatives occurred with both `NewHybrids` (2 out of 20) and `STRUCTURE` (6 out of 20) for microsatellite data. On the other hand, `NewHybrids` and `STRUCTURE` analyses of simulated RADseq data were 100% accurate in inferring ancestry (*Fig. 3B*, *Fig. S15*).

### Phylogenetic approaches clarify relationships within *murinus*

A `SplitsTree NeighborNet` phylogenetic network (*Fig. 4A*) showed a very clear separation between *griseorufus* and *murinus* with little phylogenetic conflict, and strong intraspecific structure in *murinus*, thus agreeing with previous analysis by Weisrock et al. (2010). The three well-defined clades within *murinus* correspond to the three predefined populations (western *mur*-W, contact zone area *mur*-C, and eastern *mur*-E), and in accordance with geographical distances, *mur*-W appears to be the most divergent *murinus* population. Similarly, *griseorufus* samples clustered by population (western *gri*-W and contact zone area *gri*-C), but the clades were less well-defined than in *murinus*. The only notable, though still minor, interspecific phylogenetic conflict was observed along the edges between *murinus* and *rufus* (*Fig. 4A*). All putative hybrids fell squarely within one of the two clades, with individual assignments in perfect agreement with clustering approaches. Similarly, a `NeighborNet` network using only contact zone individuals showed little to no phylogenetic conflict (*Fig. S16*).

**Fig. 4:**
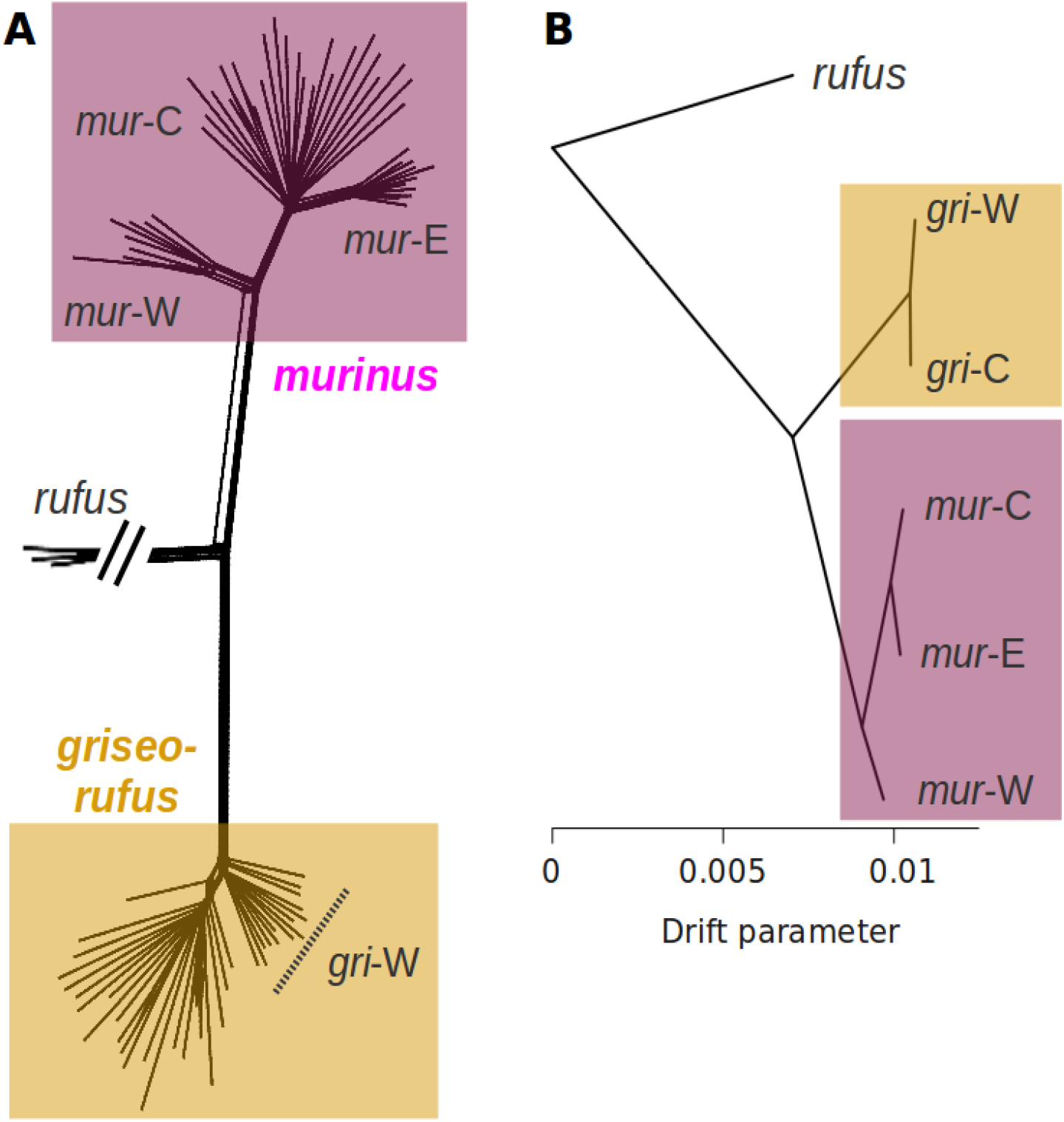
Phylogenetic relationships. **A)** A `SplitsTree` NeighborNet phylogenetic network. Each tip represents an individual, and the width of any edge boxes depicts phylogenetic conflict, which can be due to incomplete lineage sorting or admixture. Very little conflict is observed along the edges between *griseorufus* and *murinus*. *Murinus* is separated into three clades which correspond to western (*mur*-W), contact zone area (*mur*-C), and eastern (*mur*-E) populations. The separation of *griseorufus* into clades corresponding to western (*gri*-W) and contact zone area (*gri*-C) populations is not as well-defined. **B)** `Treemix` results with no migration edges. `Treemix` supports the relationships suggested by the phylogenetic network, with western *murinus* (*mur*-W) being the most divergent among the three *murinus* populations.

`Treemix` (*Fig. 4B*) was run with *murinus* and *griseorufus* individuals assigned to the five populations and *M. rufus* as the outgroup, and confirmed the relationships within *murinus* suggested by `Splitstree`: *mur*-W was the most divergent and *mur*-C and *mur*-E were sister. No significant migration edges were found between *murinus* and *griseorufus,* with instead several significant edges between *M. rufus* and *griseorufus* and *M. rufus* and *murinus* (*Fig. S17*). When *M. rufus* was excluded, significant migration edges between *griseorufus* and *murinus* did emerge, but did not include any between contact zone area populations (*gri*-C and *mur*-C) (*Fig. S18*).

### No current – but some ancestral – interspecific gene flow

D-statistics showed an over-representation of shared derived sites between both *griseorufus* populations (*gri*-W and *gri*-C) and the two southeastern *murinus* populations (*mur*-C and *mur*-E; relative to their sister *mur*-W, western *murinus*) (*Fig. 5A*). Values of D were highly similar regardless of which of the *griseorufus* or southeastern *murinus* populations were used, which suggests historical admixture between the ancestral *griseorufus* and southeastern *murinus* lineages, as well as a lack of ongoing gene flow in the contact zone. A lack of ongoing gene flow was further supported by values of D very close to (and not significantly different from) zero for comparisons testing for excess derived allele sharing between contact zone populations of both species relative to their sister populations (*Fig. 5A*).

**Fig. 5:**
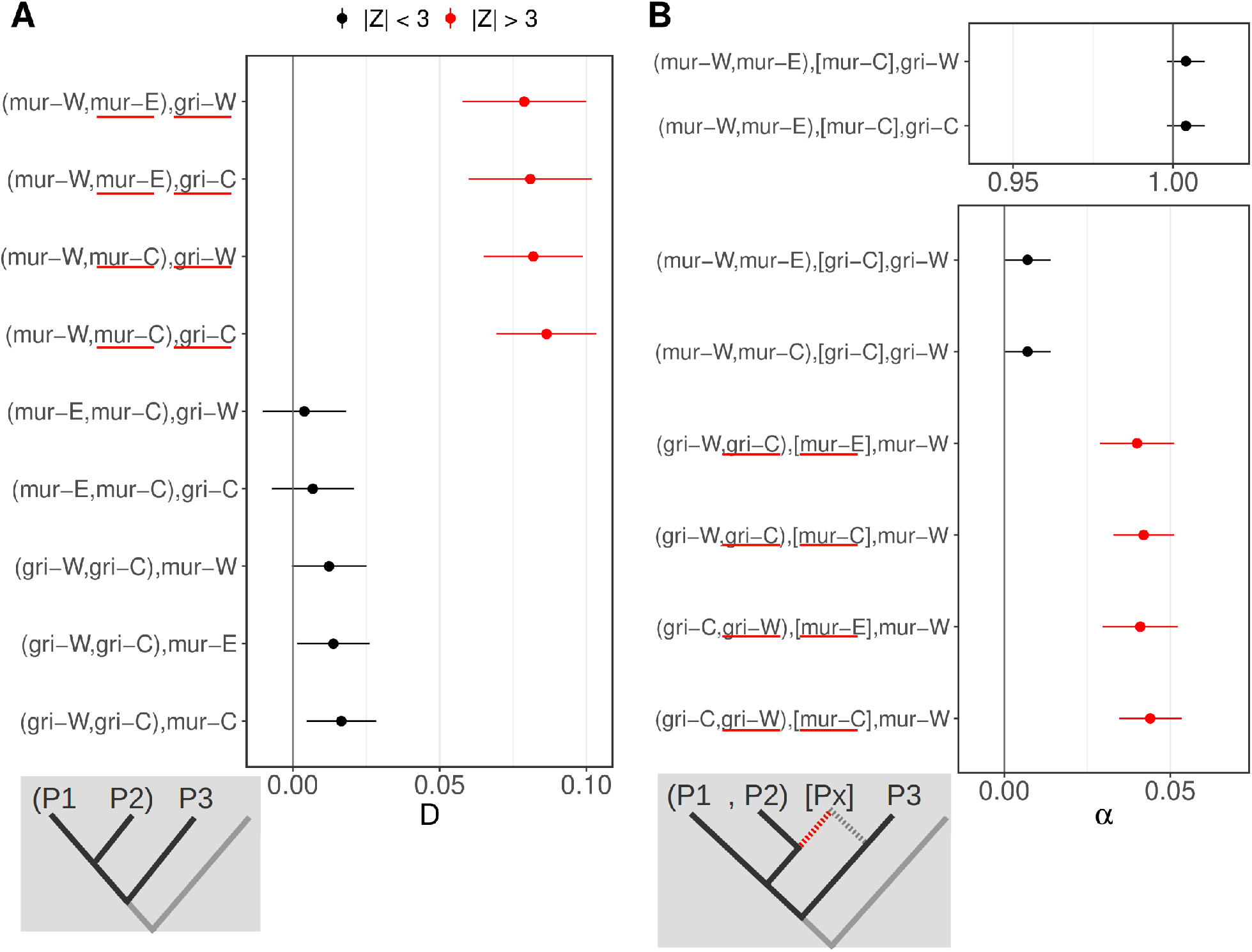
Admixture statistics suggest some ancestral but no contemporary gene flow. **A) D-statistics.** Focal comparisons are listed as `(P1,P2),P3` and test for admixture between P3 and P1 (negative D) or P2 (positive D). Populations inferred to have experiences admixture are underlined in red. For all tests, *M. rufus* was used as the outgroup (O/P4). In the top 4 rows, with *mur*-W as P1, D is significant and highly similar regardless of which *griseorufus* population (*gri*-W or *gri*-C) is used as P3 and regardless of which southeastern *murinus* population (*mur*-E or *mur*-C) is used as P2. This suggests historical but no ongoing admixture between the ancestral *griseorufus* and southeastern *murinus* lineages. A lack of ongoing gene flow is also supported by non-significant results for the bottom five comparisons. **B) f_4_-ratio tests.** Focal comparisons are listed as `(P1,P2),[Px],P3)`, where Px is tested for being a mixture between P2 and P3. On the x-axis, α indicates the proportion of P2 ancestry in Px (α=1 if Px is sister to P2 with no admixture from P3, and α=0 if Px is sister to P3 with no admixture from P2). Admixture is inferred if α is significantly different from 0 and 1 (red dots). Consistent with results for D-statistics, admixture is inferred between the two southeastern *murinus* populations and both *griseorufus* populations, with values of α highly similar regardless of which *griseorufus* population (*gri*-W or *gri*-C) is used as P1 and which as P2, and regardless of which southeastern *murinus* population (*mur*-E or *mur*-C) is used as Px.

F_4_-ratio tests similarly indicated ancestral admixture between *griseorufus* and the ancestor of contact zone (*mur*-C) and eastern *murinus (mur*-E) populations, specifically estimating that after divergence from western *murinus*, this ancestral southeastern *murinus* population experienced about 4.0-4.4% admixture with *griseorufus* (*Fig. 5B*).

Demographic modeling using `G-PhoCS` supported the presence of non-zero but low levels of historical gene flow between ancestral southeastern *murinus* and *griseorufus* (2N_m_ = 0.02-0.03 from *griseorufus* into *murinus*, and 0.06-0.07 from *murinus* into *griseorufus*), but a lack of gene flow between extant contact zone area populations of *griseorufus* and *murinus* (2N_m_ = 0) (*Fig. 6A-B*). Furthermore, some gene flow was inferred between ancestral *murinus* (i.e., the lineage ancestral to all three sampled extant populations) and ancestral *griseorufus*, particularly from *griseorufus* into *murinus* (*2N_m_ =* 0.06-0.07). Similarly, `BPP` inferred an absence of introgression between extant populations, and some introgression between ancestral populations. But unlike `G-PhoCS, BPP` inferred that introgression occurred symmetrically, and introgression was more pronounced for the event further back in time (Fig. 6C-D; see Supplementary Results for further details).

**Fig. 6:**
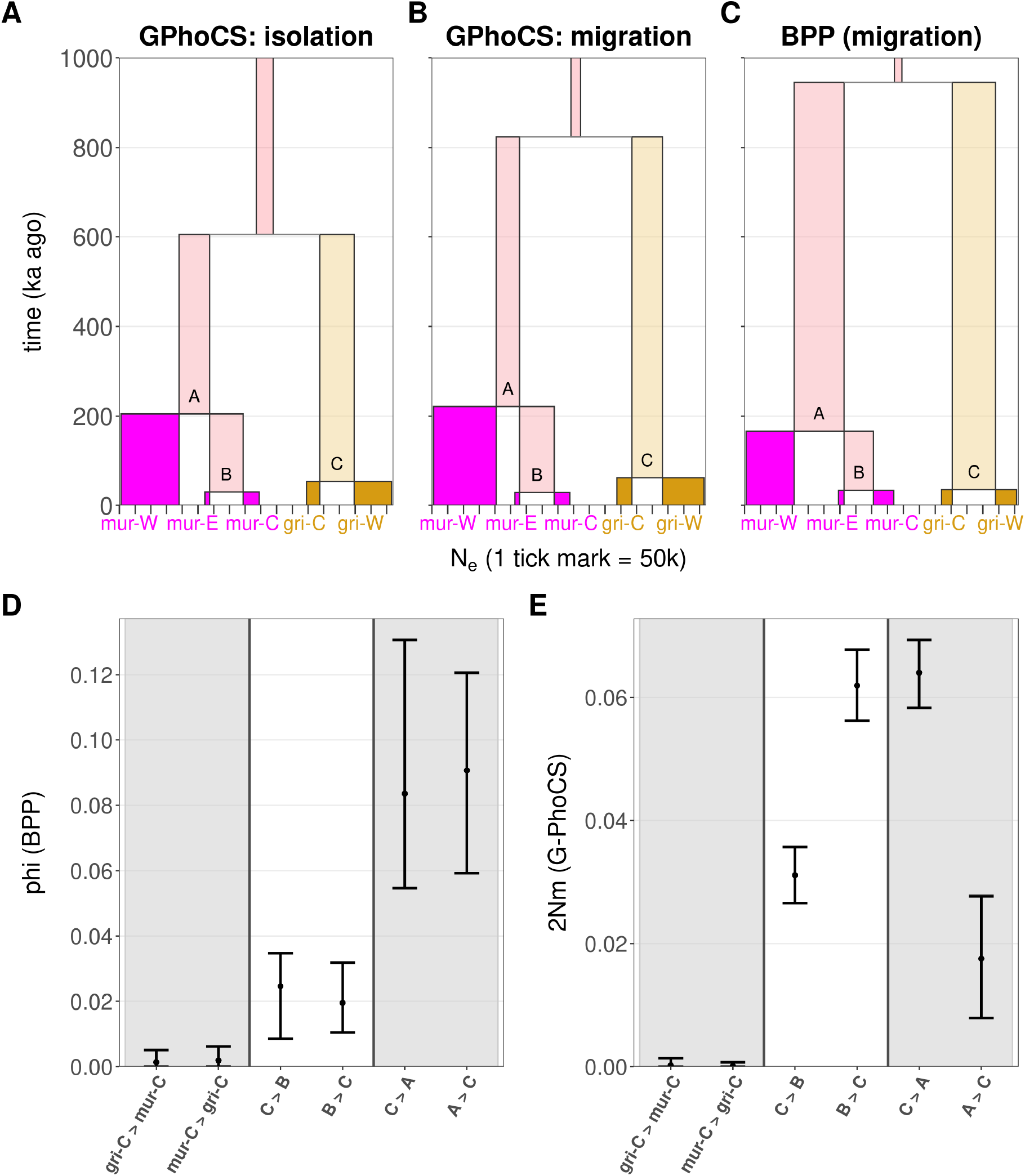
Demographic inferences using G-PhoCS and BPP. **A-C)** Summary of results for `G-PhoCS` models without (A) and with (B) gene flow and for BPP (C; with gene flow). Each box represents an extant (bright colors: gold for *griseorufus*, purple for *murinus*) or ancestral (faded colors) lineage, with box width indicating N_e_ and box height indicating time. Gene flow was estimated reciprocally between three pairs of lineages, as depicted by the arrows. **D)** Point estimates and 95% HPDs of `BPP` introgression probabilities (phi). **E)** Point estimates and 95% HPDs of `G-PhoCS` population migration rates (2Nm).

### Intraspecific differentiation is more pronounced within *murinus*

`G-PhoCS` estimated a divergence time of 20.3-37.3 ka ago (95% HPD) between the contact zone area population (*mur*-C) and eastern (*mur*-E) *murinus* populations, whereas the divergence time between western (*mur*-W) and the ancestral southeastern population (*mur*-C + *mur*-E) was inferred to be much older at 162-291 ka ago (*Fig. 6A,C*). The divergence time between western (*gri*-W) and contact zone area (*gri*-C) *griseorufus* was estimated to be 43.6-79.2 ka ago. Thus, in line with `NeighborNet` results, considerably more pronounced population structure was detected within *murinus*.

Striking differences in N_e_ between extant populations were inferred, especially in *murinus*, where those of the two southeastern populations (*mur*-E: 13-16 k, *mur*-C: 45-53k) much smaller than that of the western (*mur*-W: 194-205 k) population (*Fig. 6A,D*). Similarly, in *griseorufus*, the western (*gri*-W: 125-140 k) population was also inferred to be much larger than the southeastern population (*gri*-C: 46-50 k).

Overall, divergence time and population size estimates were similar for `G-PhoCS` models that did and those that did not incorporate gene flow (*Fig. 6C,D*) and for `BPP` (with gene flow); above, we presented estimates from `G-PhoCS` models that did incorporate gene flow. The largest differences were found for the divergence time between *murinus* and *griseorufus*, which was estimated to be 605 (95% HPD: 432-782) ka ago by `G-PhoCS` without accounting for gene flow, 824 (601-1081) ka ago by `G-PhoCS` when accounting for gene flow, and 945 (679-1238) ka ago by `BPP`.

## Discussion

We re-examined a contact zone between two species of mouse lemur in southeastern Madagascar, where extensive hybridization had previously been reported based primarily on evidence from microsatellite data (Hapke et al. 2011). With RADseq data, we found no evidence for the presence of admixed individuals, and using simulations and re-analyses of microsatellite data, we showed that previously detected hybrids were likely false positives. By including allopatric populations and performing multispecies coalescent analyses, we furthermore found a general lack of ongoing gene flow, and very low levels of ancestral gene flow, between these two species. We discuss the implications for speciation in mouse lemurs, and for inferring hybridization using microsatellites.

### Reconciling the lack of evidence for hybrids with microsatellite results

We found no admixed nuclear ancestry in any of the individuals from the contact zone. Our data is expected to have high power in species assignment and hybrid detection, given the combination of the relatively high number of genetic markers used (Vähä and Primmer 2006; McFarlane and Pemberton 2019) and the pronounced genetic differentiation between these two species (estimated divergence time in a no-migration scenario: ~600 ka ago, *Fig. 6*; average F_ST_ in the contact zone area: 0.40, *Table S5*). Furthermore, in a re-analysis of microsatellite data using the same methods as the original studies (Hapke et al. 2011, Lüdemann 2018), though restricted to the individuals used in this study, all but one of the previously detected hybrids were no longer classified as such (*Fig. 3A*).

Considering the clear and robust RADseq results, it is highly unlikely that true hybrids were missed in our analyses, even with the more limited sampling of individuals used in this study. Instead, our results suggest specifically that the hybrids inferred in Hapke et al. (2011) were false positives, and more generally, that the inference of hybridization using microsatellites can be sensitive to such false positives, particularly when using the program `NewHybrids`. While simulations showed an overall much lower accuracy of ancestry inference with microsatellites compared to SNPs, `STRUCTURE` suffered from false negatives only, whereas `NewHybrids` produced 4 false positives among 40 simulated unadmixed individuals (*Fig. 3B*). Additionally, in our reanalysis of the microsatellite data, the single individual that `NewHybrids` continued to assign hybrid ancestry to did not show signs of admixture using `STRUCTURE` (*Fig. 3A*). In Hapke et al. (2011, their Figure 5), `STRUCTURE` did also not consistently infer admixed ancestry for several of the putative hybrids. This was especially apparent when parapatric populations were included, in which case only 4 out of the 12 `NewHybrids` positives showed admixed ancestry using `STRUCTURE` (and 3 out of those 4 were still assigned <10% admixed ancestry by `STRUCTURE`, Hapke et al. 2011, their Figure 5). Even though `NewHybrids` appears considerably more prone to false positives than `STRUCTURE`, the latter did show admixed ancestry for 7 individuals in an analysis using only individuals from the contact zone site Mangatsiaka (versus 9 with `NewHybrids`).

### Ancestral gene flow and the possibility of geographically restricted gene flow

Consistent with the lack of evidence for admixed individuals in contact zone sites, we found a lack of evidence for ongoing gene flow using multiple methods, including a phylogenetic network (Fig. 4A), `Treemix` (Fig. 4B), formal admixture statistics (Fig. 5) and two multispecies coalescent methods (`G-PhoCS` and `BPP`, Fig. 6). The latter two methods did indicate some ancestral gene flow between *griseorufus* and the southeastern *murinus* populations as a whole, though these coalescent-based methods also suggested gene flow that is even more ancient (prior to population divergence within *murinus*).

In this study, we exclusively used samples from the area studied by Hapke et al. (2011), while the area examined by Gligor et al. (2009), who also inferred hybridization using microsatellites, is located 40 km further south. Based on the results of this study, what can we say about the possibility that hybridization is in fact taking place in that area, given that our coalescent analyses did not detect ongoing gene flow between *griseorufus* and *murinus*? First, the *murinus* population reported to hybridize in Gligor et al. (2009) likely involves a differentiated *Microcebus* population that has recently been split as *M. manitatra* based on patterns of genetic differentiation (Hotaling et al. 2016). Therefore, it is possible that local gene flow in that area remained undetected by our analyses, particularly when occurring from *griseorufus* into *murinus* / *M. manitatra.* However, unaccounted-for genetic differentiation between *murinus* populations may have also impacted the analyses in Gligor et al. (2009), given that their three “reference” populations likely included populations from both of the two recent splits *M. manitatra* and *M. ganzhorni* (Hotaling et al. 2016).

We also note that Gligor et al. (2009) used the same 9 microsatellite loci as Hapke et al. (2011) and applied similar analytical methods, although they used `GeneClass` rather than `NewHybrids`. Furthermore, concordance between `STRUCTURE` and `GeneClass` analyses were low (see their Fig. 5). Finally, Gligor et al. (2009) found some evidence that the ecotone populations may form their own cluster. Given the historical isolation at very small scales identified in this region, it is thus feasible that the “ecotone” population has also been isolated, further complicating ancestry inference. All in all, a genomic study using samples from that area is needed to clarify whether hybridization is taking place in the contact zone area studies by Gligor et al. (2009).

### Lack of ongoing gene flow and implications for speciation

The presence of at least two individuals with mitonuclear discordance (a *griseorufus*-type mitochondrial haplotype, and *murinus* nuclear DNA) may suggest some ongoing or recent gene flow between the two species. However, we did not detect gene flow between extant *murinus* and *griseorufus* populations using formal admixture statistics (*Fig. 4*) or with coalescent-based demographic modeling (*Fig. 5*). Combined with the lack of evidence for nuclear admixture in the contact zone, and syntopic occurrence at least one of the contact zone sites (*Fig. 2*), these findings strongly suggest that *murinus* and *griseorufus* are currently reproductively isolated.

Little is known about the relative importance of different types of reproductive isolation in mouse lemurs. Across their ranges, *murinus* and *griseorufus* occur in distinct habitat types, with *griseorufus* mostly limited to spiny forests that appear to be too arid for *murinus* (Yoder et al. 2002; Rakotondranary and Ganzhorn 2011; Rakotondranary et al. 2011a). Separation by habitat (e.g., Wuesthoff et al. 2021) at larger scales could therefore minimize or even prevent syntopic co-occurrence despite nominal sympatry in the contact zone area, thus limiting interactions between the species. At one of the two sympatric sites included in this study, Tsimelahy, species-specific sampling locations are indeed consistent with separation by habitat, at Mangatsiaka, the two species co-occur even at a very fine spatial scale but despite statistical microhabitat and dietary separation (Rakotondranary and Ganzhorn 2011; Rakotondranary et al. 2011b; *Fig. 2C*). Therefore, the observed lack of gene flow is unlikely to simply be a by-product of separation by habitat, and additional sources of pre- and/or postzygotic reproductive isolation need to be invoked.

One potential source of prezygotic reproductive isolation may be related to differences in torpor patterns: *murinus* seems to enter torpor more frequently than *griseorufus* prior to the reproductive period (Rakotondranary and Ganzhorn 2011). In several other cases of mouse lemur sympatry, differences in body size and seasonal timing of reproduction have also been observed (Evasoa et al. 2018). However, the size difference between *murinus* and *griseorufus* is modest, with the former being on average about 10-15% heavier, while it is not presently known whether timing of reproduction differs among sympatric populations (Rakotondranary et al. 2011a). Although divergence times are relatively short, postzygotic incompatibilities may also play a role. More research into sources of reproductive isolation among these and other mouse lemur species is clearly needed.

We estimated the divergence time between these two species to be less than 1 million years ago (Fig. 6). Similarly, a recent study estimated that two pairs of sympatric mouse lemur species in northeastern Madagascar each diverged less than 1 ma ago (Poelstra et al. 2020). These findings tentatively suggest an *upper bound* for the time to completion of speciation in mouse lemurs of under a million years. By comparison, Curnoe et al. (2006) found that the median estimated divergence time between pairs of naturally hybridizing primate species was 2.9 Ma. These divergence time comparisons are, however, not straightforward: dates for most other primate clades were calculated using fossil-calibrated relaxed clock methods, whereas estimates from this study and Poelstra et al. (2020) are based on coalescent analyses using mutation rates estimated from pedigree studies Poelstra et al. (2020) found large differences between these two types of divergence time estimates for the mouse lemur TMRCA (see also Tiley et al. 2020). Moreover, we here used the recent mouse lemur mutation rate estimate from Campbell et al. (2021)), whereas Poelstra et al. (2020) used a mean primate mutation rate that was 19% lower, leading to relatively older absolute time estimates.

Concerns have been raised that mouse lemurs may have undergone oversplitting or “taxonomic inflation” (Tattersall 2007; Markolf et al. 2011). The evidence for relatively rapid speciation discussed above suggests that despite limited genetic differentiation between recently described species, such species may in fact be partially or even fully reproductively isolated. On the other hand, our results suggest that the current taxonomic treatment of *M. murinus* is not tenable. Hotaling et al. (2016) recently split two southeastern micro-endemics from *M. murinus.* We estimate that the divergence time between one of these (*M. ganzhorni*) and another southeastern *murinus* population is as recent as ~40 ka ago (*Fig. 6*). Moreover, divergence between the “western” and “southeastern” population groups was much more ancient, such that *murinus s.s.* is currently paraphyletic. To fully re-evaluate the taxonomy of *murinus s.l.*., a study is needed that also includes samples from the other recent split, *M. manitatra*, and a broader sampling of western *murinus* (which itself has been shown to contain phylogeographic breaks – Pastorini et al. 2003; Schneider et al. 2010),

As per the results of our study, there are as yet no well-documented hybrid zones between mouse lemur species. This is noteworthy given the high diversity of the genus in a relatively small area. On the other hand, and perhaps even more strikingly, there are also relatively few instances of overlapping ranges between mouse lemurs. More generally, factors that limit the tempo of a successful transition of allopatrically speciating lineages into sympatry include interactions between incipient species after contact (e.g., reproductive isolation and competitive exclusion) and processes that limit such contact in the first place (e.g., low dispersal distances). Many mouse lemur species have spatially abutting ranges that are separated by large rivers, which are thought to provide barriers to dispersal for many Malagasy micro-endemics, including mouse lemurs (Martin 1972; Pastorini et al. 2003; Goodman and Ganzhorn 2004; Wilmé et al. 2006; Olivieri et al. 2007). These observations, in combination with the inference of relatively rapid evolution of reproductive isolation (in this study and in Poelstra et al. (2020)) lead us to speculate that dispersal may be a key limiting factor for generating alpha diversity in mouse lemurs.

### Contrasting and parallel demographic patterns

Intraspecific genetic differentiation was found to be considerably more pronounced in *murinus* than in *griseorufus*. To some extent, this is not surprising given the large gap in the distribution of *murinus* in southern Madagascar (*Fig. 1*). Indeed, the deepest split within *murinus* corresponds to differentiation between populations on opposite sides of this large geographic gap, and we estimated the divergence time between these populations to be over 200 ka ago (*Fig. 6*). Perhaps more striking is that differentiation between *murinus* populations within southeastern Madagascar, only ~35 km apart, is similar to that between *griseorufus* from southeastern and southwestern Madagascar, ~275 km apart (*Fig. 3A*, *Table S5*). This might be taken to suggest differences in, for example, dispersal distances between the two species. Yet, in comparing sympatric and parapatric sites within the contact zone area (*Fig. 1*), we found slightly stronger population structure within *griseorufus* (*Fig. 2*, *Fig. S12*). Therefore, general differences in dispersal patterns between *murinus* and *griseorufus* may not underlie the contrasting patterns of intraspecific differentiation at larger scales. Instead, stronger genetic differentiation in *murinus* may reflect a greater degree of historical isolation of mesic compared to more arid habitats during the Pleistocene, such as the isolation of mesic mountain areas (home to *M. manitatra*) from drier lowland sites during colder periods (Wilmé et al. 2006), or reductions and expansions of the eastern littoral forests during associated fluctuations of the sea level (Virah-Sawmy et al. 2009).

For both species, we found large and parallel differences in N_e_ between extant populations: smaller population sizes were inferred in eastern than in western populations (*Fig. 5*). Moreover, very similar effective population sizes were inferred for contact zone populations of each species (*mur*-C and *gri*-C, *Fig. 5*). The overall magnitude of intraspecific differences in N_e_ was larger in *murinus*, with a more than 10-fold difference between the western (*mur*-W) and the southeastern Mandena population (*mur*-E). The inferred small N_e_ for the Mandena population (see also Montero et al. 2019) is consistent with this population’s habitat: littoral forests are the Madagascar’s smallest and most endangered forest ecosystem (Ganzhorn et al. 2001; Virah-Sawmy et al. 2009). Previous studies, which assumed that a narrow coastal strip along the entire eastern coast originally consisted of littoral forest, estimated that ~90% of littoral forests have disappeared due to anthropogenic deforestation (Ganzhorn et al. 2001; Consiglio et al. 2006). More recent studies suggest that the forest was naturally fragmented and interspersed by heathlands, at least during the past 6,000 years, and thus prior to human arrival (Virah-Sawmy 2009; Virah-Sawmy et al. 2009).

## Conclusions

Using RADseq data, we found no evidence for admixture between two species of mouse lemurs in a contact zone in southern Madagascar. This is in sharp contrast to a previous study that found widespread hybridization among the same samples using microsatellites. Our results suggest that the hybrids inferred by the previous study were likely false positives, and we urge caution when using microsatellites to infer hybridization. Finally, we used coalescent models to show that despite an estimated divergence time of under 1 million years between these two species, interspecific gene flow only took place between ancestral populations and has long ago ceased towards the present.

## Acknowledgments

We would like to thank Ryan Campbell, Peter Larsen, and Kelsie Hunnicutt for their help with the planning of the RADseq, Anne Veillet for help with RADseq library preparation, and Tobias L. Lenz for comments on an earlier version of the manuscript.

Financial support has been provided by the Deutsche Forschungsgemeinschaft (DFG Ga 342/19), the Landesforschungsförderung Hamburg, and an Award of the Humboldt Foundation to ADY.

Field work has been carried out under the Accord de Collaboration between Madagascar National Parks, the University of Antananarivo and the University of Hamburg.

## Data Availability

Raw sequence data will be made available through the NCBI. Processed data, such as VCF and FASTA files, will be made available through the Dryad Digital Repository. All code used to run the analyses and produce the figures in this manuscript can be found on GitHub at https://github.com/jelmerp/lemurs_contactzone_grimur.

