## Supplementary Materials for "RADseq data reveal a lack of admixture in a mouse lemur contact zone contrary to previous microsatellite results"

### 1 Supplementary Methods

#### 1.1 Genotype filtering

We used `GATK`'s "`HaplotypeCaller`" tool to produce GVCF files for each sample. GVCF files were then merged to multi-sample, single-scaffold "*GenomicsDB Workspaces*" using `GATK`'s "`GenmicsDBImport`" tool (this merging step is necessary because in GATK v4 since the joint genotyping tool n longer accepts multiple GVCF files), and these *Workspaces* were then used as input for `GATK`'s "`GenotypeGVCFs`" tool, which we ran using the "`--use-new-qual-calculator`" option.

VCF files were filtered according to recommendations from O'Leary et al. (2018), following their "FS6" filtering steps (See Table 2 in O'Leary et al. 2018). First, we selected only SNPs (i.e., discarding structural variants) using `GATK`'s `SelectVariants` tool with option "`-selectType SNP`". Next, we annotated the VCF file with the *Allele Balance* statistic using `GATK's` "`VariantAnnotator`" tool with option "`-A AlleleBalance`". The FS6 filtering procedure involved removing the following data in the following order:

1. Per-sample genotypes with a depth below 5 (using "`--minDP 5`" in `vcftools`) are are set to "missing".
2. Sites with an across-sample genotype quality (*Qual*) lower than 20 (using "`--minQual 20`" in `vcftools`).
3. Sites with an across-sample mean depth lower than 15 (using "`--min-meanDP 15`" in `vcftools`).
4. Sites/samples with specified amounts of missing data. A progressive series of filtering steps for missing data was performed, alternating between filtering at the site level (wherein sites with more than the specified percentage of missing data are removed) and filtering at the sample level (wherein sites with more than the specified percentage of missing data are removed). Site-filtering was done using "`--max-missing <threshold>`" in `vcftools`, whereas sample-filtering first required the computation of per-sample statistics on missing data (using "`--missing-indv`" in `vcftools`), followed by the removal of samples exceeding the threshold using "`--remove <sample_ID>`" in `vcftools`. Specifically, these are the maximum missing data thresholds for the consecutive filtering

steps (note that this procedure is the same as in O'Leary et al.'s FS6, but the following notation is slightly different):

a) site: 50%

b) sample: 90%

c) site: 40%

d) sample: 70%

e) site: 30%

f) sample: 50%

5. Sites with more than two alleles. Note that this step is not present in O'Leary's FS6 filter.
6. Sites not passing any of the following filters based on statistics in the "INFO" field of the VCF file.

a) Allele balance (*ABHet* < 0.2 or *ABHet* > 0.8)

b) Mapping quality ratio of reference vs. alternative allele (*MQRankSum* < -12.5)

c) Strandedness of reference vs. alternative allele (*FS* > 60.0)

d) Quality-by-depth ratio (*QD* < 2.0)

e) Absolute mapping quality (*MQ* < 40.0)

f) Read position of reference vs. alternative allele (*ReadPosRankSum* < -8)

These sites were set to "*FILTER*" status using `GATK`'s "`VariantFiltration`" tool followed by removal using "`--remove-filtered-all`" in `vcftools`. The threshold values used here are based on GATK guidelines (https://gatkforums.broadinstitute.org/gatk/discussion/2806/howto-apply-hard-filters-to-a-call-set), except for item a, the allele balance filter, which is not present in the GATK recommendations. For this step, we largely followed the GATK guidelines because O'Leary et al. (2018) were not always explicit about thresholds and/or performed filtering with custom procedures rather than "`VariantFiltration`" parameters. Additional minor differences with O'Leary's et al. (2018) framework are that we also included filtering based on absolute mapping score (item e above) and read position (item f above), and that we removed non-properly paired reads during BAM-file processing steps rather than in the VCF.

7. Sites with a depth higher than the mean depth plus twice the standard deviation in depth. This maximum depth threshold was computed in a first step, and sites with excessive depth were next removed using "`–max-meanDP <threshold>`" in `vcftools`.

8. A final round of filtering sites and samples with excessive missing data (as in step 5, see above) were removed using:

    a) sample: 25%

    b) site: 5%

For analyses on subsets of the data, such as the 6-species and 4-species datasets used in the `G-PhoCS` analysis, we started by subsetting the *raw* VCF with the focal samples, and then applying the filters as listed above.

For the set of samples from the contact zone area, we additionally produced two datasets using more lenient filtering procedures, to be able to examine admixture using more samples and SNPs:

1. A dataset produced by omitting the last round of removal of SNPs and samples based on missing data (i.e., step 8 above). This is referred to in the supplementary figures and tables below as "***FS7***".

2. A dataset produced using the FS6 filter without the sample-level filtering steps (in steps 4 and 8). Sample selection was instead done by taking all samples that passed the regular FS6 filter, and additionally retaining two additional putative hybrids and two samples with mitonuclear discordance with reasonable depth of coverage. This is referred to in the supplementary figures and tables below as "***rescue***".

#### 1.2 Creating full-sequence FASTA files

To produce full-sequence FASTA files from the VCF files (i.e., including invariant sites), we used the following procedure: First, we ran the `GATK` v3.8 tool "`FastaAlternateReferenceMaker`" for each sample, producing a single-sample whole-genome fasta file based on the reference genome but replacing sites that were called as non-reference alleles in the *unfiltered* VCF file. Then, the following classes of bases were masked: (1) non-reference bases that did not pass filtering (see RADseq genotyping and filtering section above); (2) sites that were classified as non-callable using `GATK` v3.8's "`CallableLoci`" tool, using a minimum depth of 3.

Next, "loci" were extracted by (1) defining loci at an individual level as stretches of at least 25 consecutive called (non-N) bases or multiple such stretches separated by at most 10 consecutive Ns; (2) intersecting these stretches across samples, requiring at least 1 bp of overlap, and trimming from both ends any bases with fewer than 80% of focal samples represented; (3) filtering the resulting loci to include only loci that: (a) had sequences for at least 90% of focal samples, (b) were at least a 100 bp long, and (c) contained at most 10% missing data.

#### 1.3 Ancestry assignment analyses with Structure

`STRUCTURE` performs a clustering of samples into a user defined number of clusters K, based on the genotype. The result are posterior probabilities for each sample belonging to each of the K clusters. For this analysis, no prior population information was provided, using the admixture ancestry model with independent allele frequencies and lambda set to 1. As suggested by Gilbert et al. (2012), 20 runs per K were performed, with 100 000 iterations of burn-in and 100 000 iterations after that, while K was set from 1 to 5. Selecting for the optimal number of K was done by following the Evanno et al. (2005) method of the ad hoc statistic $\Delta K$, and the Pritchard et al. (2000) method, both implemented in the web version of `Structure Harvester` version v0.6.94 (Earl and vonHoldt 2012). For every dataset, $\Delta K$ clearly indicated $K = 2$ as best K, whereas the Pritchard method suggests a $K = 4$.

## 1.4 $F_{ST}$

Weir and Cockerham weighted $F_{ST}$ estimates for pairwise comparisons between population groupings was calculated from VCF files using `vcftools`.

### 2 Supplementary Tables

**Table S1: Sample information: contact zone area**

“put_hybrid” = putative hybrid, “sp” = species, “mgri” = *Microcebus griseorufus*, “mmur” = *Microcebus murinus*, “pop” = population designation.

| ID | Sample_ID | Hapke_ID | site | pop | poptype | lat | lon | sp_radseq | sp_msat | sp_mtDNA | filter_pass | put_hybrid |
|---|---|---|---|---|---|---|---|---|---|---|---|---|
| mgri068 | 886 | NA | Mangatsiaka | gri-C | sympatric | -24.963083 | 46.555761 | mgri | mgri | mmur | fail | 1 |
| mgri069 | 887 | NA | Mangatsiaka | gri-C | sympatric | -24.963147 | 46.557383 | mgri | mgri | mmur | rescue | 1 |
| mgri070 | 889 | NA | Mangatsiaka | gri-C | sympatric | -24.963244 | 46.557797 | mgri | mgri | mgri | FS7 | 0 |
| mgri071 | 890 | NA | Mangatsiaka | gri-C | sympatric | -24.962222 | 46.557378 | mgri | mgri | mmur | rescue | 1 |
| mgri072 | 2019 | NA | Mangatsiaka | gri-C | sympatric | -24.962831 | 46.557511 | mgri | mgri | mgri | FS7 | 0 |
| mgri073 | 2020 | NA | Mangatsiaka | gri-C | sympatric | -24.962650 | 46.557983 | mgri | mgri | mgri | fail | 0 |
| mgri074 | 2021 | NA | Mangatsiaka | gri-C | sympatric | -24.962219 | 46.557972 | mgri | mgri | mgri | fail | 0 |
| mgri075 | HZ03 | Hzf_Mg_Hamb032 | Hazofotsy | gri-C | parapatric | -24.841575 | 46.528875 | mgri | mgri | mgri | FS6 | 0 |
| mgri076 | HZ05 | Hzf_Mg_Hamb034 | Hazofotsy | gri-C | parapatric | -24.828919 | 46.547633 | mgri | mgri | mgri | FS6 | 0 |
| mgri077 | HZ06 | Hzf_Mg_Hamb035 | Hazofotsy | gri-C | parapatric | -24.828011 | 46.549208 | mgri | mgri | mgri | FS6 | 0 |
| mgri078 | HZ11 | Hzf_Mg_Hamb040 | Hazofotsy | gri-C | parapatric | -24.841658 | 46.529031 | mgri | mgri | mgri | FS6 | 0 |
| mgri079 | HZ12 | Hzf_Mg_Hamb041 | Hazofotsy | gri-C | parapatric | -24.841658 | 46.529031 | mgri | mgri | mgri | FS6 | 0 |
| mgri080 | HZ13 | Hzf_Mg_Hamb042 | Hazofotsy | gri-C | parapatric | -24.842269 | 46.529461 | mgri | mgri | mgri | FS6 | 0 |
| mgri081 | HZ15 | Hzf_Mg_Hamb104 | Hazofotsy | gri-C | parapatric | -24.841350 | 46.528883 | mgri | mgri | mgri | fail | 0 |
| mgri082 | HZ16 | Hzf_Mg_Hamb105 | Hazofotsy | gri-C | parapatric | -24.841350 | 46.528883 | mgri | mgri | mgri | FS6 | 0 |
| mgri083 | MG05 | Mtk_Mg_Hamb059 | Mangatsiaka | gri-C | sympatric | -24.964172 | 46.557636 | mgri | mgri | mgri | fail | 0 |
| mgri084 | MG06 | Mtk_Mg_Hamb060 | Mangatsiaka | gri-C | sympatric | -24.966453 | 46.554600 | mgri | mgri | mgri | FS6 | 0 |
| mgri085 | MG16 | Mtk_Mg_Hamb126 | Mangatsiaka | gri-C | sympatric | -24.966244 | 46.554728 | mgri | mgri | mgri | FS6 | 0 |
| mgri086 | MG20 | Mtk_Mg_Hamb130 | Mangatsiaka | gri-C | sympatric | -24.962811 | 46.556350 | mgri | mgri | mgri | FS6 | 0 |
| mgri087 | MG22 | Mtk_Mg_Hamb132 | Mangatsiaka | gri-C | sympatric | -24.966244 | 46.554728 | mgri | mgri | mgri | FS6 | 0 |
| mgri088 | MG25 | Mtk_Mg_Hamb135 | Mangatsiaka | gri-C | sympatric | -24.964172 | 46.557636 | mgri | mgri | mgri | FS6 | 0 |
| mgri089 | MG85 | NA | Mangatsiaka | gri-C | sympatric | -24.962850 | 46.558133 | mgri | mgri | mgri | fail | 0 |
| mgri090 | Micro13 | Tml_Mg_MAndo13 | Tsimelahy | gri-C | sympatric | -24.951069 | 46.617919 | mgri | mgri | mgri | FS6 | 0 |
| mgri091 | Micro14 | Tml_Mg_MAndo14 | Tsimelahy | gri-C | sympatric | -24.956069 | 46.610750 | mgri | mgri | mgri | fail | 0 |
| mgri092 | Micro19 | Tml_Mg_MAndo19 | Tsimelahy | gri-C | sympatric | -24.955589 | 46.613069 | mgri | mgri | mgri | fail | 0 |
| mgri093 | Micro31 | Tml_Mg_MAndo31 | Tsimelahy | gri-C | sympatric | -24.955589 | 46.613069 | mgri | mgri | mgri | FS6 | 0 |
| mgri094 | Micro44 | Tml_Mg_MAndo44 | Tsimelahy | gri-C | sympatric | -24.952181 | 46.616931 | mgri | mgri | mgri | FS6 | 0 |
| mgri095 | Micro58 | Tml_Mg_MAndo58 | Tsimelahy | gri-C | sympatric | -24.956150 | 46.610481 | mgri | mgri | mgri | FS6 | 0 |
| mgri096 | Micro60 | Tml_Mg_MAndo60 | Tsimelahy | gri-C | sympatric | -24.951100 | 46.617739 | mgri | mgri | mgri | FS7 | 0 |
| mgri097 | Micro70 | Tml_Mg_MAndo70 | Tsimelahy | gri-C | sympatric | -24.955931 | 46.611169 | mgri | mgri | mgri | FS6 | 0 |
| mgri098 | Micro81 | Mtk_Mg_MAndo81 | Mangatsiaka | gri-C | sympatric | -24.968836 | 46.557314 | mgri | mgri | mgri | FS6 | 0 |
| mgri099 | TM41 | Tml_Mg_Hamb078 | Tsimelahy | gri-C | sympatric | -24.956050 | 46.610808 | mgri | mgri | mgri | FS6 | 0 |
| mgri100 | TM46 | Tml_Mg_Hamb082 | Tsimelahy | gri-C | sympatric | -24.955550 | 46.612750 | mgri | mgri | mgri | FS6 | 0 |

| ID | Sample_ID | Hapke_ID | site | pop | poptype | lat | lon | sp_radseq | sp_msat | sp_mtDNA | filter_pass | put_hybrid |
|---|---|---|---|---|---|---|---|---|---|---|---|---|
| mgri101 | TM49 | Tml_Mg_Hamb085 | Tsimelahy | gri-C | sympatric | -24.955578 | 46.612522 | mgri | mgri | mgri | FS7 | 0 |
| mgri102 | TM50 | Tml_Mg_Hamb086 | Tsimelahy | gri-C | sympatric | -24.955822 | 46.613642 | mgri | mgri | mgri | FS7 | 0 |
| mgri103 | TM51 | Tml_Mg_Hamb087 | Tsimelahy | gri-C | sympatric | -24.956211 | 46.616492 | mgri | mgri | mgri | FS6 | 0 |
| mgri104 | TM52 | Tml_Mg_Hamb088 | Tsimelahy | gri-C | sympatric | -24.956325 | 46.615497 | mgri | mgri | mgri | FS6 | 0 |
| mhyb001 | 2006 | NA | Mangatsiaka | gri-C | sympatric | -24.963147 | 46.557383 | mgri | hybrid | mgri | rescue | 1 |
| mhyb002 | 2009 | NA | Mangatsiaka | mur-C | sympatric | -24.962558 | 46.555864 | mmur | hybrid | mmur | rescue | 1 |
| mhyb003 | 2026 | NA | Mangatsiaka | gri-C | sympatric | -24.963083 | 46.555761 | mgri | hybrid | mgri | FS6 | 1 |
| mhyb004 | 2027 | NA | Mangatsiaka | gri-C | sympatric | -24.963339 | 46.557489 | mgri | mgri | mgri | FS6 | 0 |
| mhyb005 | 2033 | NA | Mangatsiaka | gri-C | sympatric | -24.963339 | 46.557489 | mgri | hybrid | mgri | fail | 1 |
| mhyb006 | 2034 | NA | Mangatsiaka | gri-C | sympatric | -24.962981 | 46.557908 | mgri | hybrid | mgri | FS6 | 1 |
| mhyb007 | MG03 | NA | Mangatsiaka | mur-C | sympatric | -24.962461 | 46.553428 | mmur | hybrid | mmur | FS6 | 1 |
| mhyb008 | MG15 | Mtk_Mm_Hamb125 | Mangatsiaka | mur-C | sympatric | -24.966886 | 46.554289 | mmur | hybrid | mmur | FS6 | 1 |
| mhyb009 | MG18 | Mtk_Mm_Hamb128 | Mangatsiaka | mur-C | sympatric | -24.960756 | 46.560169 | mmur | hybrid | mmur | FS6 | 1 |
| mhyb010 | MG24 | Mtk_Mm_Hamb134 | Mangatsiaka | mur-C | sympatric | -24.963300 | 46.555264 | mmur | hybrid | mmur | FS6 | 1 |
| mhyb011 | MG26 | Mtk_Mg_Hamb136 | Mangatsiaka | gri-C | sympatric | -24.962131 | 46.558203 | mgri | hybrid | mgri | FS6 | 1 |
| mhyb012 | MG27 | Mtk_Mm_Hamb137 | Mangatsiaka | mur-C | sympatric | -24.963703 | 46.554400 | mmur | hybrid | mmur | FS6 | 1 |
| mhyb013 | MG34 | Mtk_Mm_Hamb144 | Mangatsiaka | mur-C | sympatric | -24.962181 | 46.553358 | mmur | hybrid | mmur | FS6 | 1 |
| mhyb014 | MG50 | Mtk_Mm_Hamb160 | Mangatsiaka | mur-C | sympatric | -24.963556 | 46.553606 | mmur | hybrid | mmur | FS6 | 1 |
| mhyb015 | MG64 | Mtk_Mg_Hamb167 | Mangatsiaka | gri-C | sympatric | -24.962983 | 46.557056 | mgri | hybrid | mgri | FS6 | 1 |
| mhyb016 | Micro33 | Tml_Mg_MAndo33 | Tsimelahy | gri-C | sympatric | -24.955361 | 46.613989 | mgri | hybrid | mgri | FS6 | 1 |
| mmur037 | AB01 | Abt_Mm_Hamb044 | Ambatoabo | mur-C | parapatric | -24.819172 | 46.669656 | mmur | mmur | mmur | FS6 | 0 |
| mmur038 | AB03 | Abt_Mm_Hamb046 | Ambatoabo | mur-C | parapatric | -24.819172 | 46.669656 | mmur | mmur | mmur | FS6 | 0 |
| mmur039 | AB04 | Abt_Mm_Hamb047 | Ambatoabo | mur-C | parapatric | -24.818292 | 46.671294 | mmur | mmur | mmur | FS6 | 0 |
| mmur040 | AB06 | Abt_Mm_Hamb049 | Ambatoabo | mur-C | parapatric | -24.818869 | 46.664431 | mmur | mmur | mmur | FS6 | 0 |
| mmur041 | AB12 | Abt_Mm_Hamb095 | Ambatoabo | mur-C | parapatric | -24.819028 | 46.670106 | mmur | mmur | mmur | FS6 | 0 |
| mmur042 | AB15 | Abt_Mm_Hamb098 | Ambatoabo | mur-C | parapatric | -24.818292 | 46.671294 | mmur | mmur | mmur | FS6 | 0 |
| mmur043 | AB16 | Abt_Mm_Hamb099 | Ambatoabo | mur-C | parapatric | -24.817528 | 46.672233 | mmur | mmur | mmur | FS6 | 0 |
| mmur044 | AB18 | Abt_Mm_Hamb101 | Ambatoabo | mur-C | parapatric | -24.817575 | 46.672714 | mmur | mmur | mmur | FS6 | 0 |
| mmur045 | MG17 | Mtk_Mm_Hamb127 | Mangatsiaka | mur-C | sympatric | -24.962792 | 46.558772 | mmur | mmur | mmur | FS6 | 0 |
| mmur046 | MG19 | Mtk_Mm_Hamb129 | Mangatsiaka | mur-C | sympatric | -24.963703 | 46.554400 | mmur | mmur | mmur | FS6 | 0 |
| mmur047 | MG23 | Mtk_Mm_Hamb133 | Mangatsiaka | mur-C | sympatric | -24.961067 | 46.559056 | mmur | mmur | mmur | FS6 | 0 |
| mmur048 | MG28 | Mtk_Mm_Hamb138 | Mangatsiaka | mur-C | sympatric | -24.962811 | 46.556350 | mmur | mmur | mmur | FS6 | 0 |
| mmur049 | MG42 | Mtk_Mm_Hamb152 | Mangatsiaka | mur-C | sympatric | -24.962181 | 46.553358 | mmur | mmur | mmur | FS6 | 0 |
| mmur050 | MG49 | Mtk_Mm_Hamb159 | Mangatsiaka | mur-C | sympatric | -24.964047 | 46.553792 | mmur | mmur | mmur | FS6 | 0 |
| mmur051 | MG58 | Mtk_Mm_Hamb161 | Mangatsiaka | mur-C | sympatric | -24.963556 | 46.553606 | mmur | mmur | mmur | FS6 | 0 |
| mmur052 | MG65 | Mtk_Mm_Hamb168 | Mangatsiaka | mur-C | sympatric | -24.961622 | 46.556336 | mmur | mmur | mmur | FS6 | 0 |

| ID | Sample_ID | Hapke_ID | site | pop | poptype | lat | lon | sp_radseq | sp_msat | sp_mtDNA | filter_pass | put_hybrid |
|---|---|---|---|---|---|---|---|---|---|---|---|---|
| mmur053 | MG66 | Mtk_Mm_Hamb169 | Mangatsiaka | mur-C | sympatric | -24.962247 | 46.556506 | mmur | mmur | mmur | FS6 | 0 |
| mmur054 | MG67 | Mtk_Mm_Hamb170 | Mangatsiaka | mur-C | sympatric | -24.963133 | 46.558022 | mmur | mmur | mmur | fail | 0 |
| mmur055 | MG68 | Mtk_Mm_Hamb171 | Mangatsiaka | mur-C | sympatric | -24.962797 | 46.558875 | mmur | mmur | mmur | FS6 | 0 |
| mmur056 | MG73 | Mtk_Mm_Hamb073 | Mangatsiaka | mur-C | sympatric | -24.962444 | 46.555161 | mmur | mmur | mmur | FS6 | 0 |
| mmur057 | MG75 | Mtk_Mm_Hamb075 | Mangatsiaka | mur-C | sympatric | -24.962803 | 46.553764 | mmur | mmur | mmur | FS6 | 0 |
| mmur058 | MG79 | NA | Mangatsiaka | mur-C | sympatric | -24.964111 | 46.555672 | mmur | mmur | mmur | FS6 | 0 |
| mmur059 | MG83 | NA | Mangatsiaka | mur-C | sympatric | -24.963333 | 46.554697 | mmur | mmur | mmur | FS7 | 0 |
| mmur060 | MG84 | NA | Mangatsiaka | mur-C | sympatric | -24.963978 | 46.554981 | mmur | mmur | mmur | FS7 | 0 |
| mmur061 | MG88 | NA | Mangatsiaka | mur-C | sympatric | -24.962975 | 46.553869 | mmur | mmur | mmur | FS6 | 0 |
| mmur062 | MG92 | NA | Mangatsiaka | mur-C | sympatric | -24.962753 | 46.556922 | mmur | mmur | mmur | fail | 0 |
| mmur063 | Micro04 | Tml_Mm_MAndo04 | Tsimelahy | mur-C | sympatric | -24.956250 | 46.617800 | mmur | mmur | mmur | FS6 | 0 |
| mmur064 | Micro06 | Tml_Mm_MAndo06 | Tsimelahy | mur-C | sympatric | -24.954511 | 46.620689 | mmur | mmur | mmur | FS6 | 0 |
| mmur065 | Micro12 | Tml_Mm_MAndo12 | Tsimelahy | mur-C | sympatric | -24.958350 | 46.616369 | mmur | mmur | mmur | FS6 | 0 |
| mmur066 | Micro24 | Tml_Mm_MAndo24 | Tsimelahy | mur-C | sympatric | -24.954161 | 46.619189 | mmur | mmur | mmur | FS6 | 0 |
| mmur067 | Micro25 | Tml_Mm_MAndo25 | Tsimelahy | mur-C | sympatric | -24.948131 | 46.621661 | mmur | mmur | mmur | FS6 | 0 |
| mmur068 | Micro27 | Tml_Mm_MAndo27 | Tsimelahy | mur-C | sympatric | -24.947961 | 46.621219 | mmur | mmur | mmur | FS6 | 0 |
| mmur069 | Micro38 | Tml_Mm_MAndo38 | Tsimelahy | mur-C | sympatric | -24.958350 | 46.616369 | mmur | mmur | mmur | FS6 | 0 |
| mmur070 | Micro49 | Tml_Mm_MAndo49 | Tsimelahy | mur-C | sympatric | -24.954250 | 46.619300 | mmur | mmur | mmur | FS6 | 0 |
| mmur071 | Micro88 | Mtk_Mm_MAndo88 | Mangatsiaka | mur-C | sympatric | -24.966683 | 46.554583 | mmur | mmur | mmur | FS6 | 0 |
| mmur072 | TM42 | Tml_Mm_Hamb079 | Tsimelahy | mur-C | sympatric | -24.948039 | 46.621075 | mmur | mmur | mmur | FS6 | 0 |
| mmur073 | TM43 | Tml_Mm_Hamb080 | Tsimelahy | mur-C | sympatric | -24.957525 | 46.617106 | mmur | mmur | mmur | FS6 | 0 |
| mmur074 | TM44 | Tml_Mm_Hamb081 | Tsimelahy | mur-C | sympatric | -24.954903 | 46.619461 | mmur | mmur | mmur | fail | 0 |
| mmur075 | TM47 | Tml_Mm_Hamb083 | Tsimelahy | mur-C | sympatric | -24.947964 | 46.621269 | mmur | mmur | mmur | FS7 | 0 |
| mmur076 | TM48 | Tml_Mm_Hamb084 | Tsimelahy | mur-C | sympatric | -24.948575 | 46.620556 | mmur | mmur | mmur | FS6 | 0 |
| mmur077 | TM53 | Tml_Mm_Hamb089 | Tsimelahy | mur-C | sympatric | -24.956372 | 46.617286 | mmur | mmur | mmur | FS7 | 0 |

**Table S2: Sample information: allopatric populations and outgroups**
All these samples pass the FS6 VCF filter.

“pop” = population designation.

| ID | Sample_ID | species | site | pop |
|---|---|---|---|---|
| mgan007 | 00-016A-8577 | *ganzhorni* | Mandena | mur-E |
| mgan008 | 00-016A-875E | *ganzhorni* | Mandena | mur-E |
| mgan010 | 00-016A-8A1D | *ganzhorni* | Mandena | mur-E |
| mgan011 | 00-016A-8EBC | *ganzhorni* | Mandena | mur-E |
| mgan014 | 00072854DC | *ganzhorni* | Mandena | mur-E |
| mgan016 | 0007289B19 | *ganzhorni* | Mandena | mur-E |
| mgan017 | 00074C3B6F | *ganzhorni* | Mandena | mur-E |
| mgan018 | 00074C3F7D | *ganzhorni* | Mandena | mur-E |
| mgan019 | 00074C439B | *ganzhorni* | Mandena | mur-E |
| mgan021 | 00074C54EC | *ganzhorni* | Mandena | mur-E |
| mgan022 | 00074CHC59 | *ganzhorni* | Mandena | mur-E |
| mgri005 | JMR008 | *griseorufus* | Antabore | gri-W |
| mgri006 | JMR009 | *griseorufus* | Antabore | gri-W |
| mgri007 | JMR010 | *griseorufus* | Antabore | gri-W |
| mgri008 | JMR011 | *griseorufus* | Antabore | gri-W |
| mgri037 | JMR007 | *griseorufus* | Tongaenoro | gri-W |
| mgri040 | 000611B575 | *griseorufus* | Tsimanampetsotsa | gri-W |
| mgri041 | 000611C0E8 | *griseorufus* | Tsimanampetsotsa | gri-W |
| mgri043 | 00063932CD | *griseorufus* | Tsimanampetsotsa | gri-W |
| mgri044 | 00063935DE | *griseorufus* | Tsimanampetsotsa | gri-W |
| mgri045 | 00063983C7 | *griseorufus* | Tsimanampetsotsa | gri-W |
| mgri046 | 00063999A9 | *griseorufus* | Tsimanampetsotsa | gri-W |
| mgri047 | 000639B1E7 | *griseorufus* | Tsimanampetsotsa | gri-W |
| mgri050 | JMR025 | *griseorufus* | Vombositse | gri-W |
| mgri051 | JMR026 | *griseorufus* | Vombositse | gri-W |
| mmur001 | RMR44 | *murinus* | Andranomena | mur-W |
| mmur002 | RMR45 | *murinus* | Andranomena | mur-W |
| mmur004 | RMR47 | *murinus* | Andranomena | mur-W |
| mmur006 | RMR49 | *murinus* | Andranomena | mur-W |
| mmur009 | Joerg33 | *murinus* | Kirindy | mur-W |
| mmur012 | RMR27 | *murinus* | Manamby | mur-W |
| mmur013 | RMR28 | *murinus* | Manamby | mur-W |
| mmur014 | RMR29 | *murinus* | Manamby | mur-W |
| mruf003 | RMR147 | *rufus* | Andrambovato | NA |
| mruf007 | E250M100 | *rufus* | Ranomafana | NA |
| mruf008 | E250M91 | *rufus* | Ranomafana | NA |

**Table S3: QC statistics for FASTQ, BAM, and VCF files by *microsatellite* assignment group.**

| **statistic** | ***griseorufus*** | **hybrid** | ***murinus*** |
|---|---|---|---|
| FASTQ: read length - mean | 135.5 | 135.6 | 135.6 |
| FASTQ: read length - median | 135.6 | 135.6 | 135.6 |
| FASTQ: read quality - mean | 38.24 | 38.24 | 38.24 |
| FASTQ: read quality - median | 38.25 | 38.25 | 38.24 |
| FASTQ: raw reads - mean | 8,021,390 | 6,483,790 | 6,459,171 |
| FASTQ: raw reads - median | 7,255,766 | 5,966,376 | 6,026,840 |
| FASTQ: dedupped reads - mean | 4,133,115 | 3,325,910 | 3,279,254 |
| FASTQ: dedupped reads - median | 3,668,470 | 3,044,472 | 3,077,260 |
| FASTQ: trimmed reads - mean | 3,191,330 | 2,580,940 | 2,545,889 |
| FASTQ: trimmed reads - median | 2,902,724 | 2,377,917 | 2,379,176 |
| BAM: mapped reads - mean | 2,978,376 | 2,410,410 | 2,385,964 |
| BAM: mapped reads - median | 2,689,533 | 2,230,030 | 2,242,135 |
| BAM: mapping % - mean | 93.4 % | 93.53 % | 93.87 % |
| BAM: mapping % - median | 93.38 % | 93.43 % | 93.79 % |
| BAM: properly paired % - mean | 99.76 % | 99.82 % | 99.85 % |
| BAM: properly paired % - median | 99.76 % | 99.84 % | 99.85 % |
| BAM: mapping quality - filtered - mean | 59.22 | 59.2 | 59.21 |
| BAM: mapping quality - filtered - median | 59.22 | 59.18 | 59.2 |
| BAM: mapping quality - unfiltered - mean | 44.59 | 45.22 | 45.77 |
| BAM: mapping quality - unfiltered - median | 44.61 | 45.19 | 45.77 |
| BAM: depth of coverage - mean | 25.85 | 23.38 | 24.35 |
| BAM: depth of coverage- median | 24.9 | 22.41 | 23.61 |
| VCF (FS6): depth of coverage - mean | 39.77 | 34.74 | 38.2 |
| VCF (FS6): depth of coverage - median | 38.56 | 31.96 | 35.59 |
| VCF (FS6): % missing SNPs - mean | 2.247 % | 2.917 % | 2.487 % |
| VCF (FS6): % missing SNPs - median | 1.574 % | 2.549 % | 1.894 % |

**Table S4: Filter-passing status by microsatellite assignment group.**

| **microsatellite assignment** | **standard (“FS6”)** | **last filtering step omitted (“FS7”)** | **FS6 + “rescue”** | **failed** | ***sum*** |
|---|---|---|---|---|---|
| *griseorufus* | 23 | 5 (28 total) | 2 (25 total) | 8 | 38 |
| hybrid | 12 | 0 (12 total) | 4 (18 total) | 1 | 15 |
| *murinus* | 36 | 4 (40 total) | 0 (36 total) | 3 | 43 |
| ***sum*** | 71 | 9 | 4 | 12 | 96 |

**Table S5: $F_{ST}$**

**A:** For contact zone dataset (FS6 filter).
*"sym"=sympatric, "para"=parapatric*

| **pop 1** | **pop 2** | **$F_{ST}$** |
|---|---|---|
| *griseorufus* – sym (n=21) | *murinus* – sym (n=33) | 0.398 |
| *griseorufus* – para (n=7) | *murinus* – para (n=8) | 0.419 |
| *gri*-C (sym+ para) (n=28) | *mur*-C (sym+ para) (n=41) | 0.399 |

**B:** For dataset with contact zone and allopatric populations (FS6 filter)

| | ***gri*-W** | ***mur*-C** | ***mur*-E** | ***mur*-W** |
|---|---|---|---|---|
| ***gri*-C** (n=19) | 0.060 | 0.461 | 0.504 | 0.436 |
| ***gri*-W** (n=14) | | 0.437 | 0.479 | 0.403 |
| ***mur*-C** (n=24) | | | 0.108 | 0.167 |
| ***mur*-E** (n=11) | | | | 0.243 |
| ***mur*-W** (n=8) | | | | |

### 3 Supplementary Figures

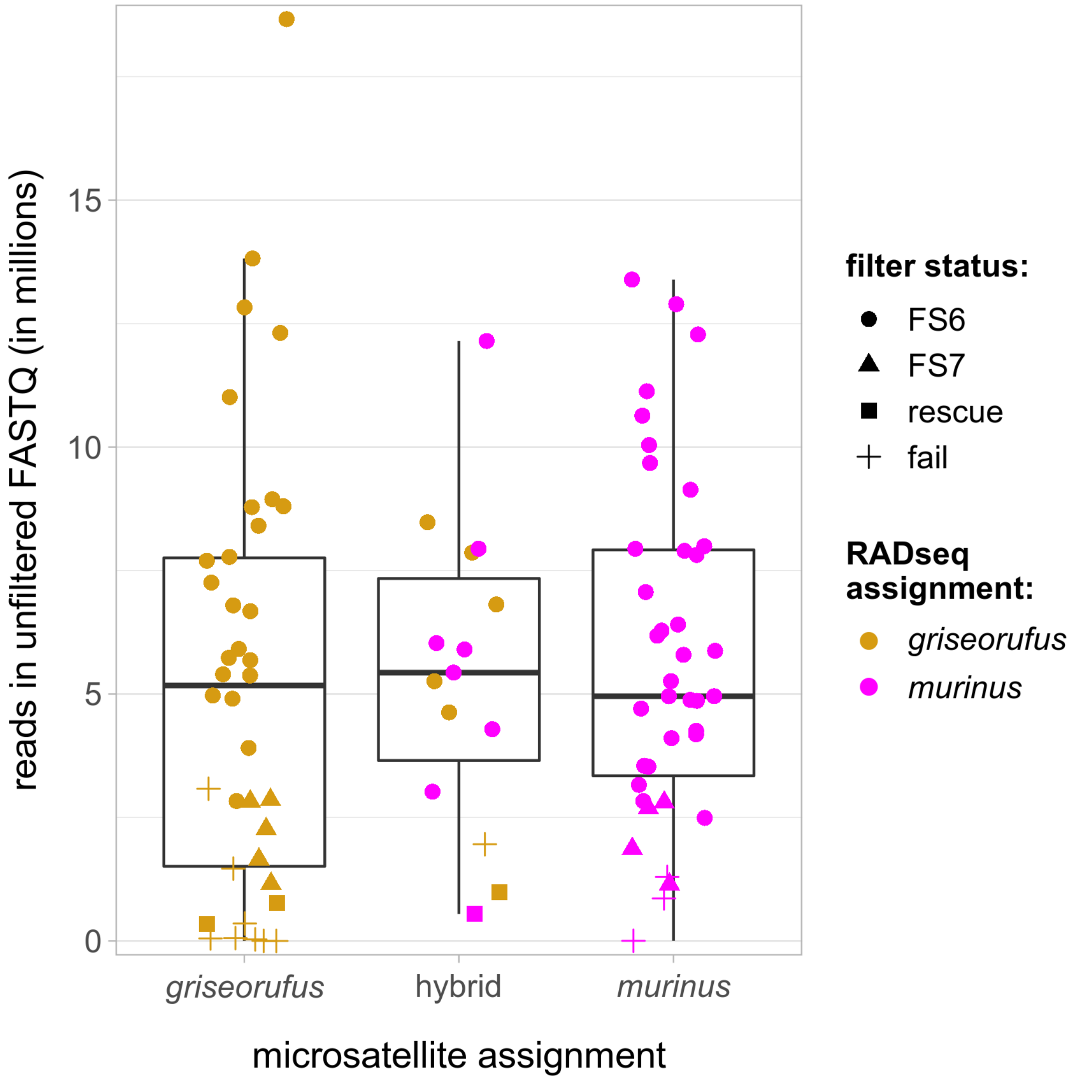

**Fig. S1: Number of reads in unfiltered FASTQ files.**
Comparison across microsatellite species assignments (x-axis), RADseq species assignments (color of jittered points), and filter status (shape of jittered points).

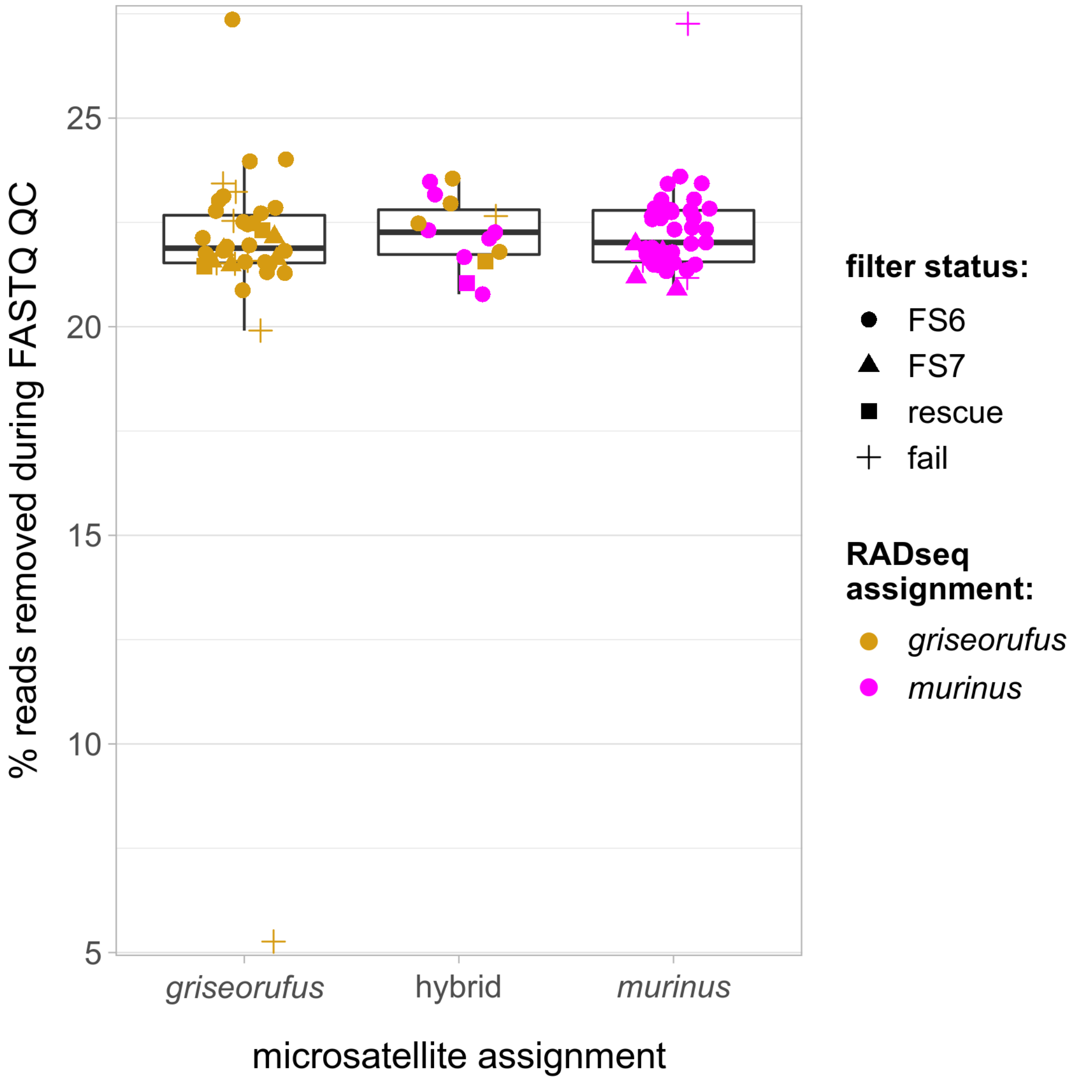

**Fig. S2: Percentage of reads removed by FASTQ filtering.**
Comparison across microsatellite species assignments (x-axis), RADseq species assignments (color of jittered points), and filter status (shape of jittered points).

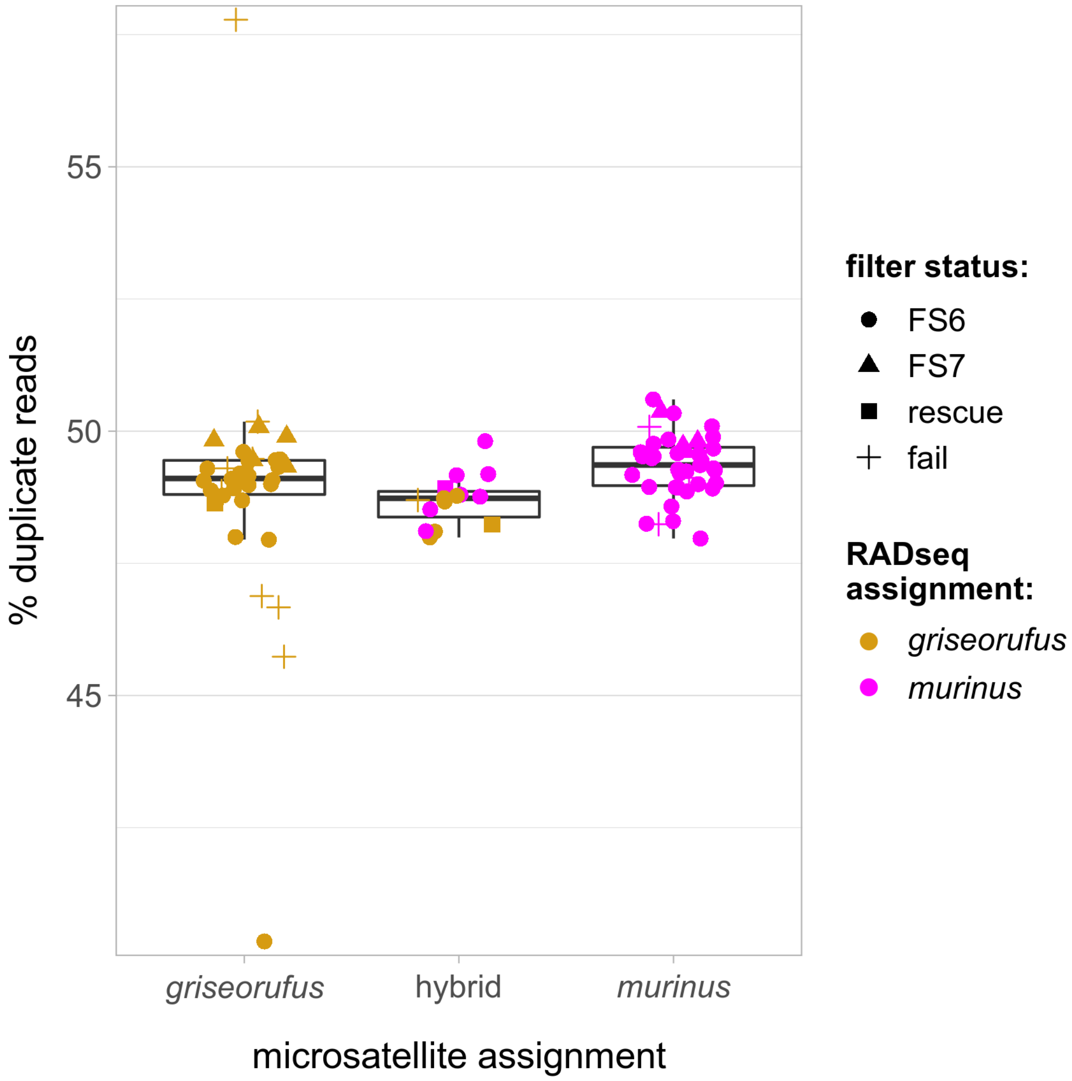

**Fig. S3: Percentage of duplicate reads.**
Comparison across microsatellite species assignments (x-axis), RADseq species assignments (color of jittered points), and filter status (shape of jittered points). Duplicate reads were removed prior to mapping.

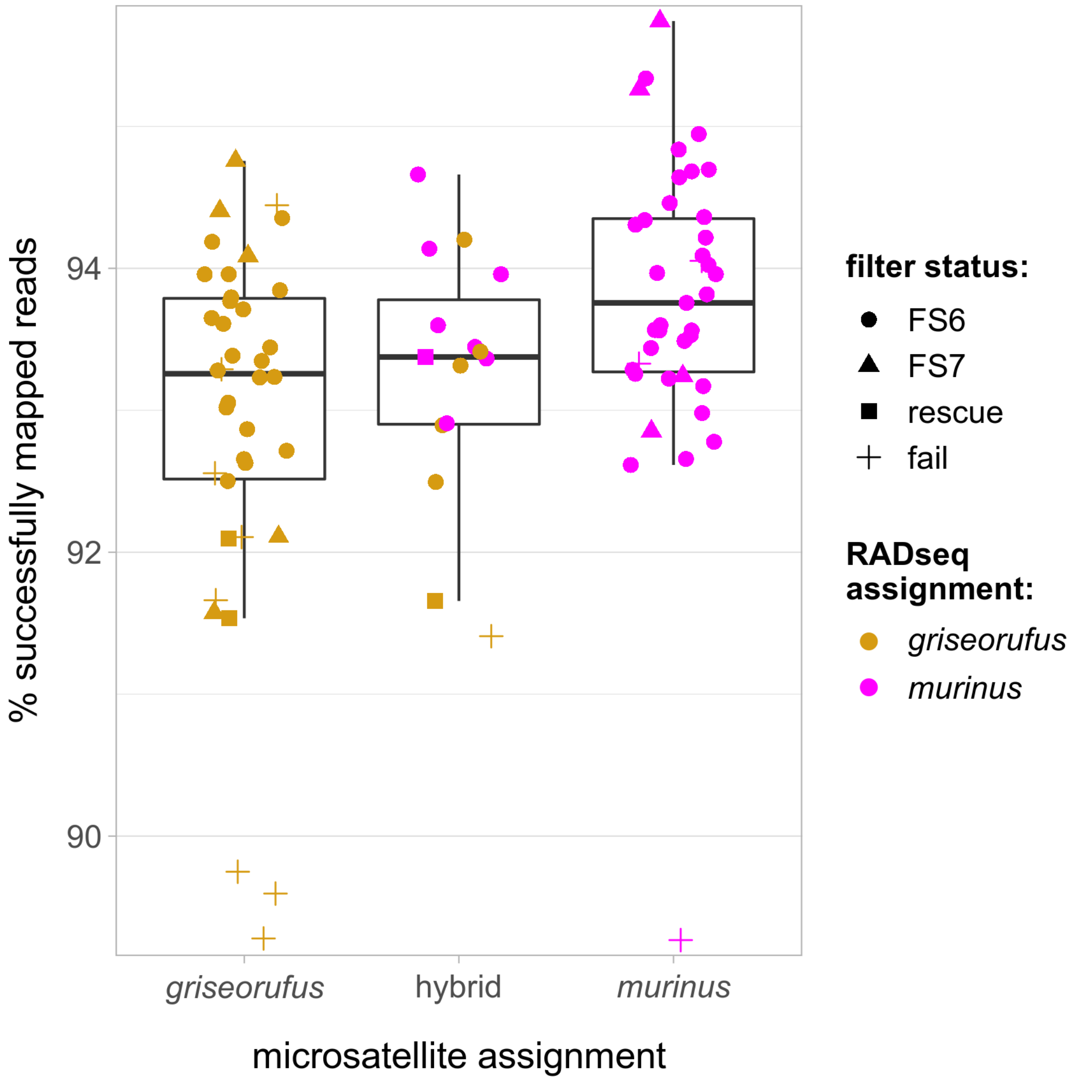

**Fig. S4: Percentage of successfully mapped reads.**
Comparison across microsatellite species assignments (x-axis), RADseq species assignments (color of jittered points), and filter status (shape of jittered points). Duplicate reads were removed prior to mapping.

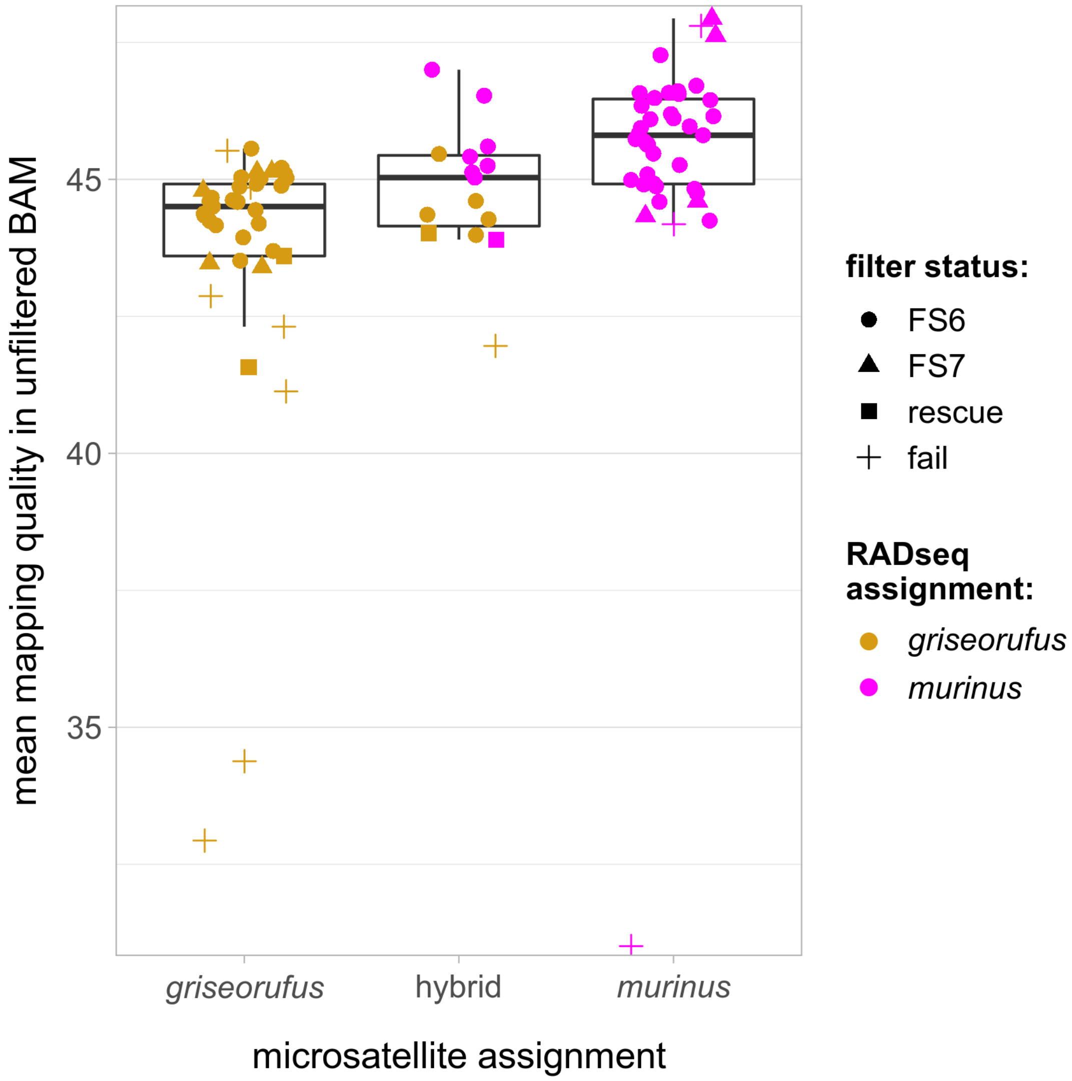

**Fig. S5: Mean mapping quality in unfiltered BAM files.**
Comparison across microsatellite species assignments (x-axis), RADseq species assignments (color of jittered points), and filter status (shape of jittered points).

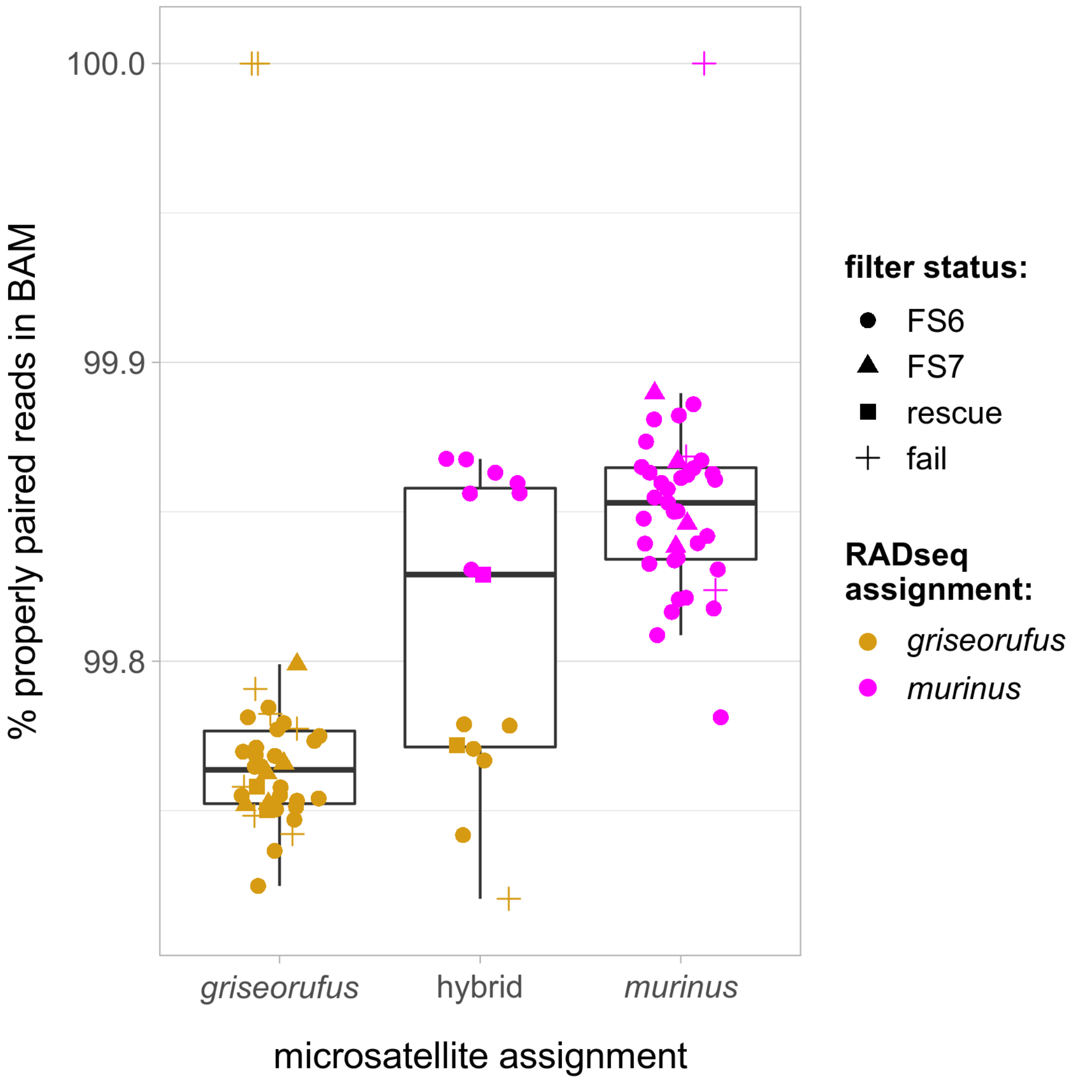

**Fig. S6: Percentage of properly paired read pairs in filtered BAM files.**
170 Comparison across microsatellite species assignments (x-axis), RADseq species assignments (color of jittered points), and filter status (shape of jittered points).

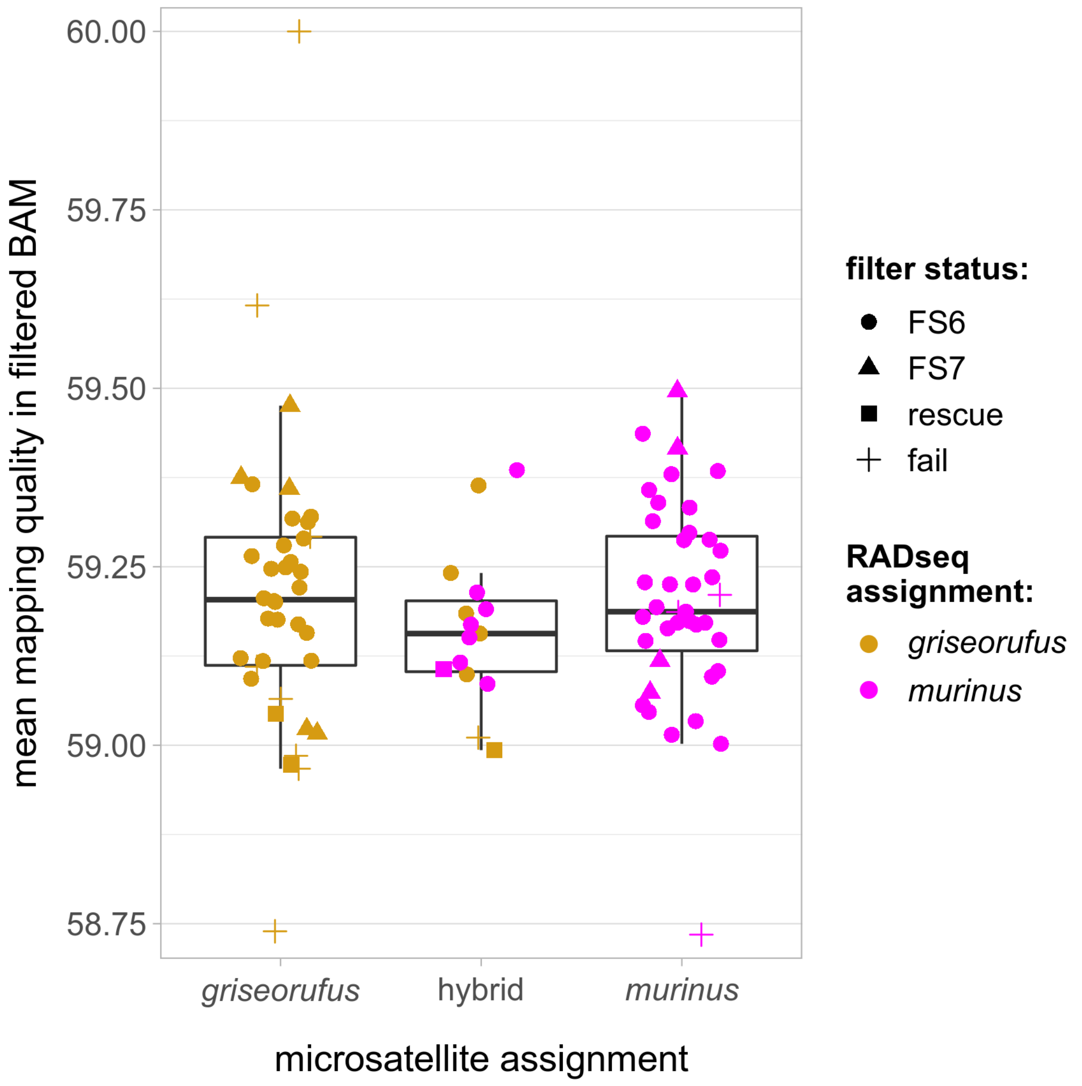

**Fig. S7: Mean mapping quality in filtered BAM files.**
Comparison across microsatellite species assignments (x-axis), RADseq species assignments (color of jittered points), and filter status (shape of jittered points).

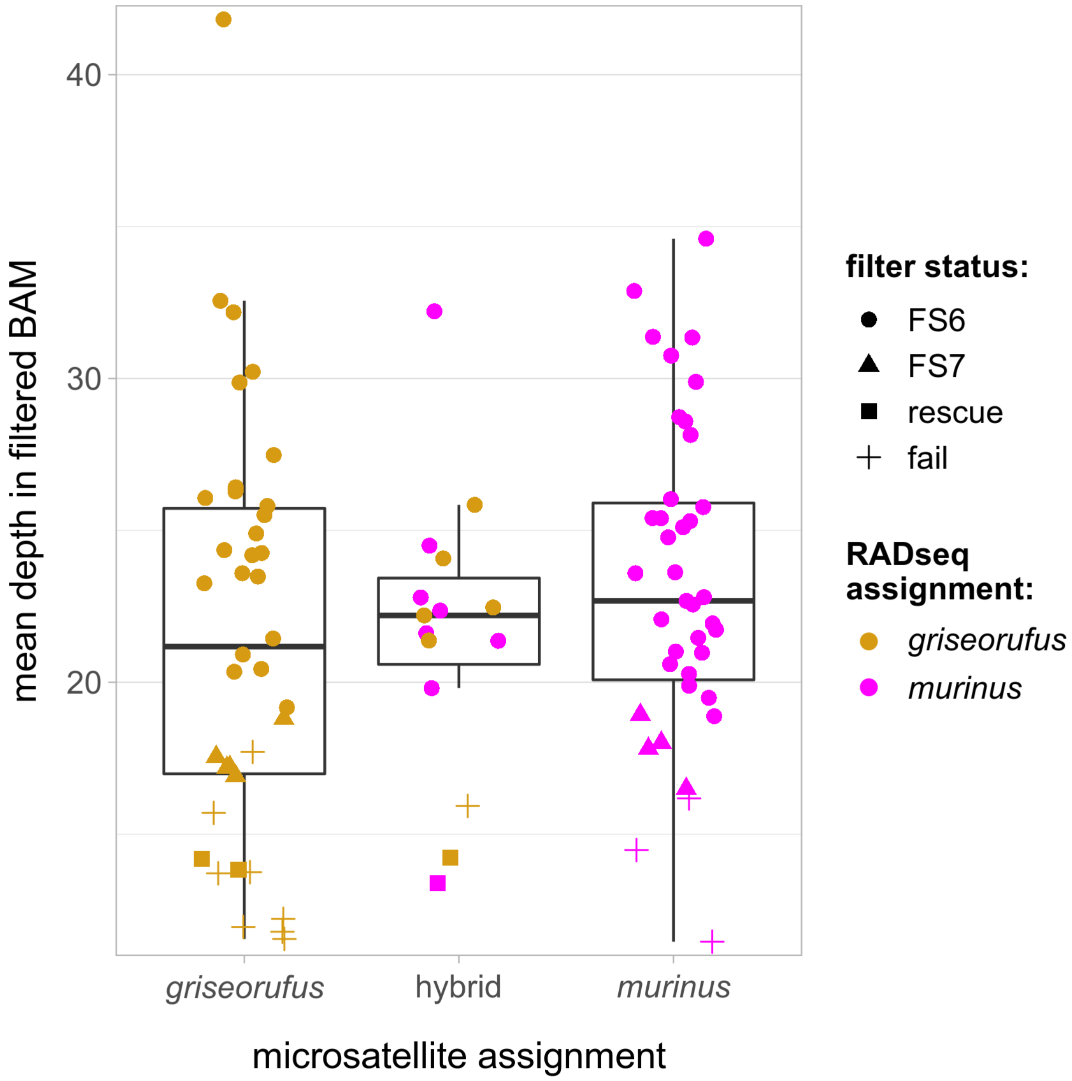

**Fig. S8: Mean depth of coverage in filtered BAM files.**
Comparison across microsatellite species assignments (x-axis), RADseq species assignments (color of jittered points), and filter status (shape of jittered points).

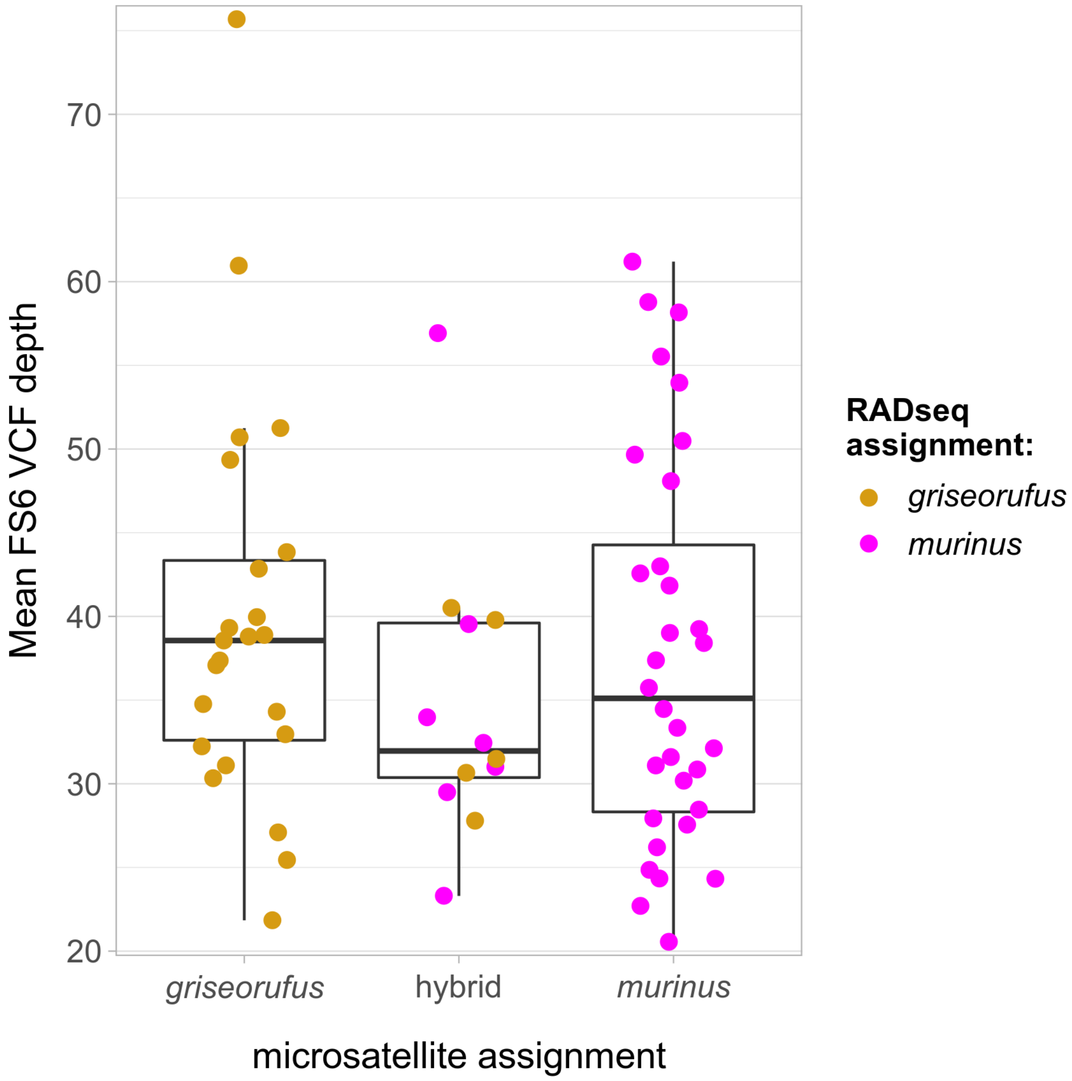

**Fig. S9: Mean depth of coverage in the "FS6" VCF file.**
Comparison across microsatellite species assignments (x-axis), RADseq species assignments (color of jittered points), and filter status (shape of jittered points). This VCF file was used for the main analyses of ancestry in the contact zone are; see Methods for details on the filtering procedure.

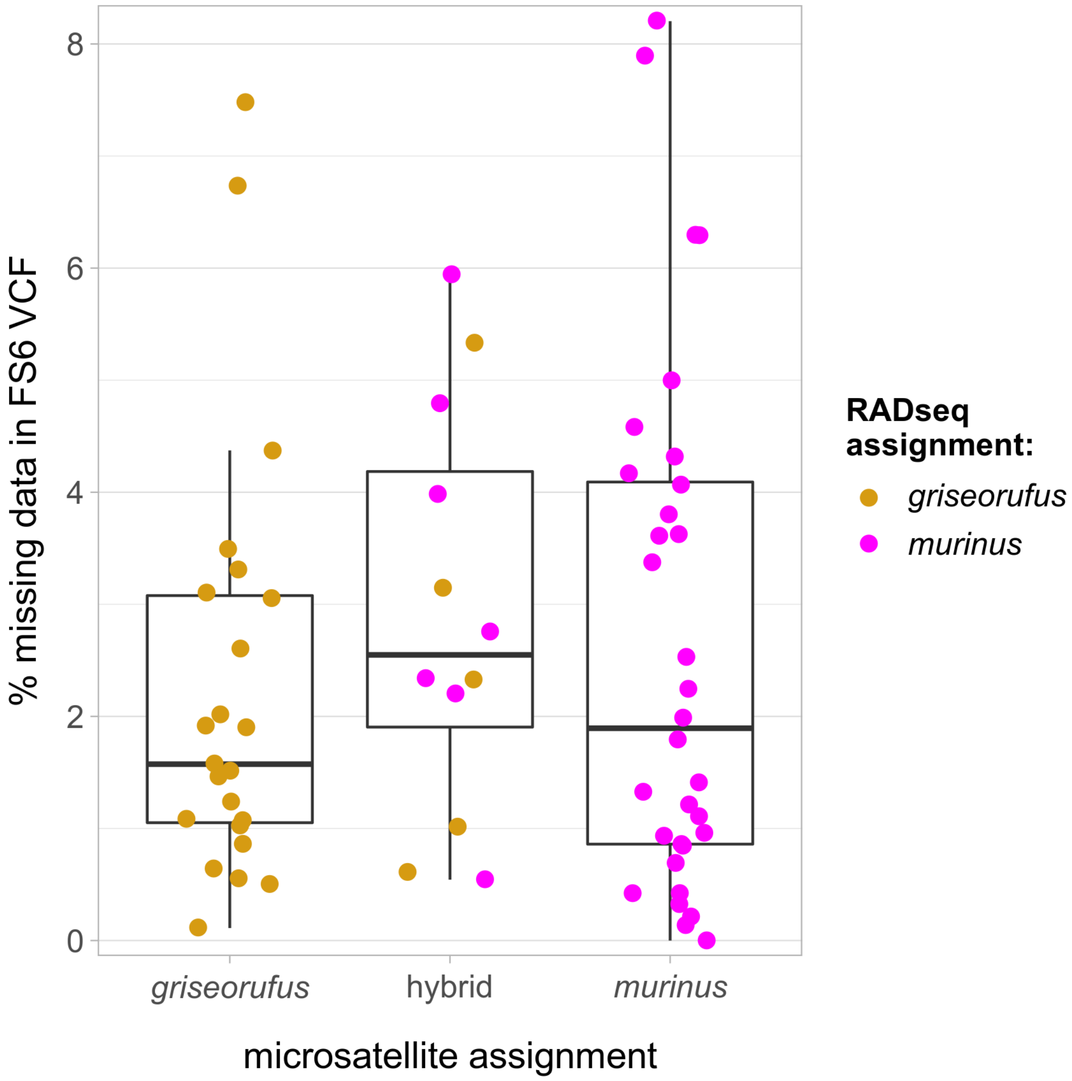

**Fig. S10: Percentage of missing data in the "FS6" VCF file.**
Comparison across microsatellite species assignments (x-axis), RADseq species assignments (color of jittered points), and filter status (shape of jittered points). This VCF file was used for the main analyses of ancestry in the contact zone are; see Methods for details on the filtering procedure.

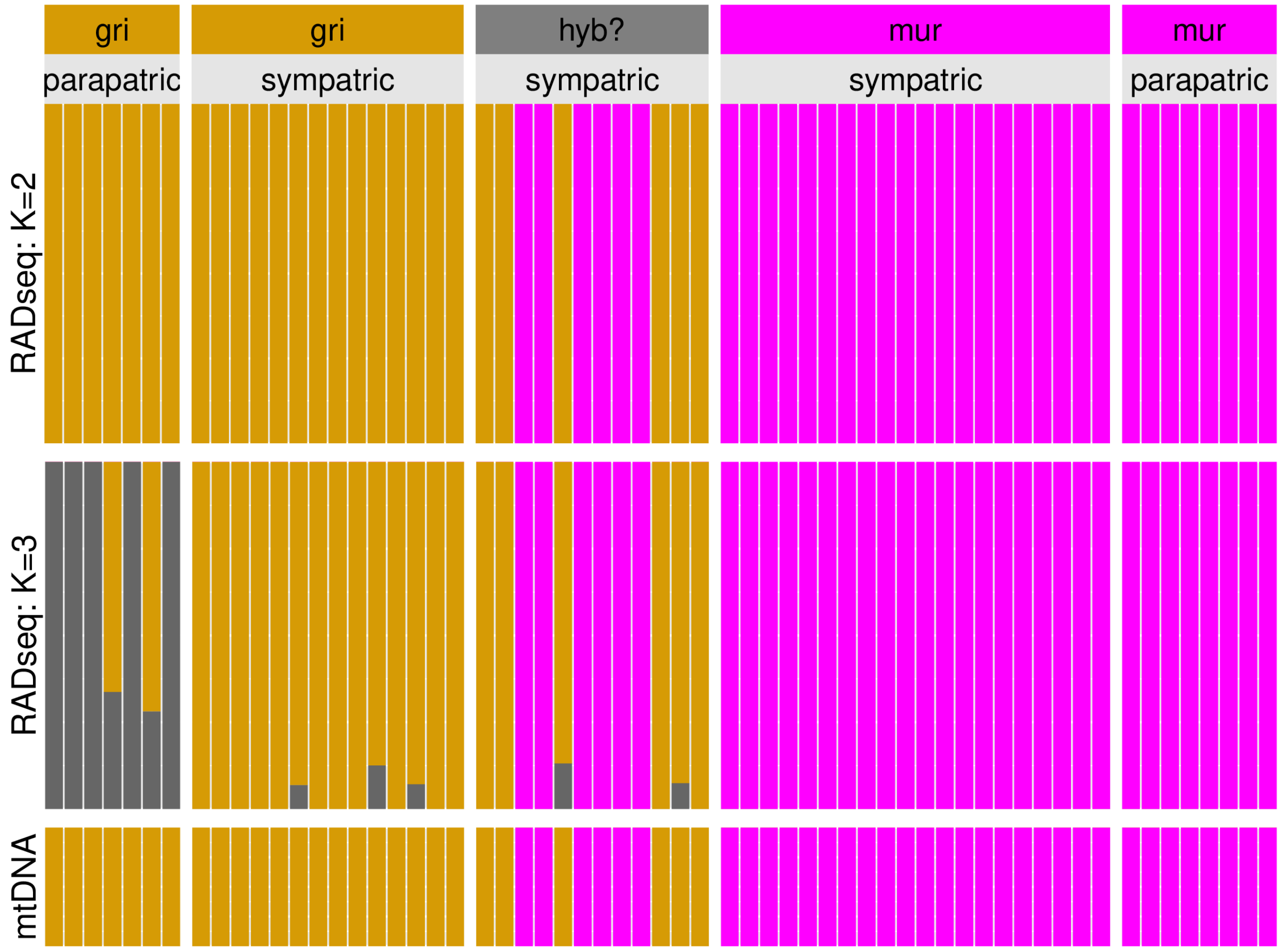

**Fig. S11: ADMIXTURE results including for K=3.**
The third cluster relates to differentiation in *griseorufus*, between parapatric (Hazofotsy) and sympatric (Mangatsiaka, Tsimelahy) sites.

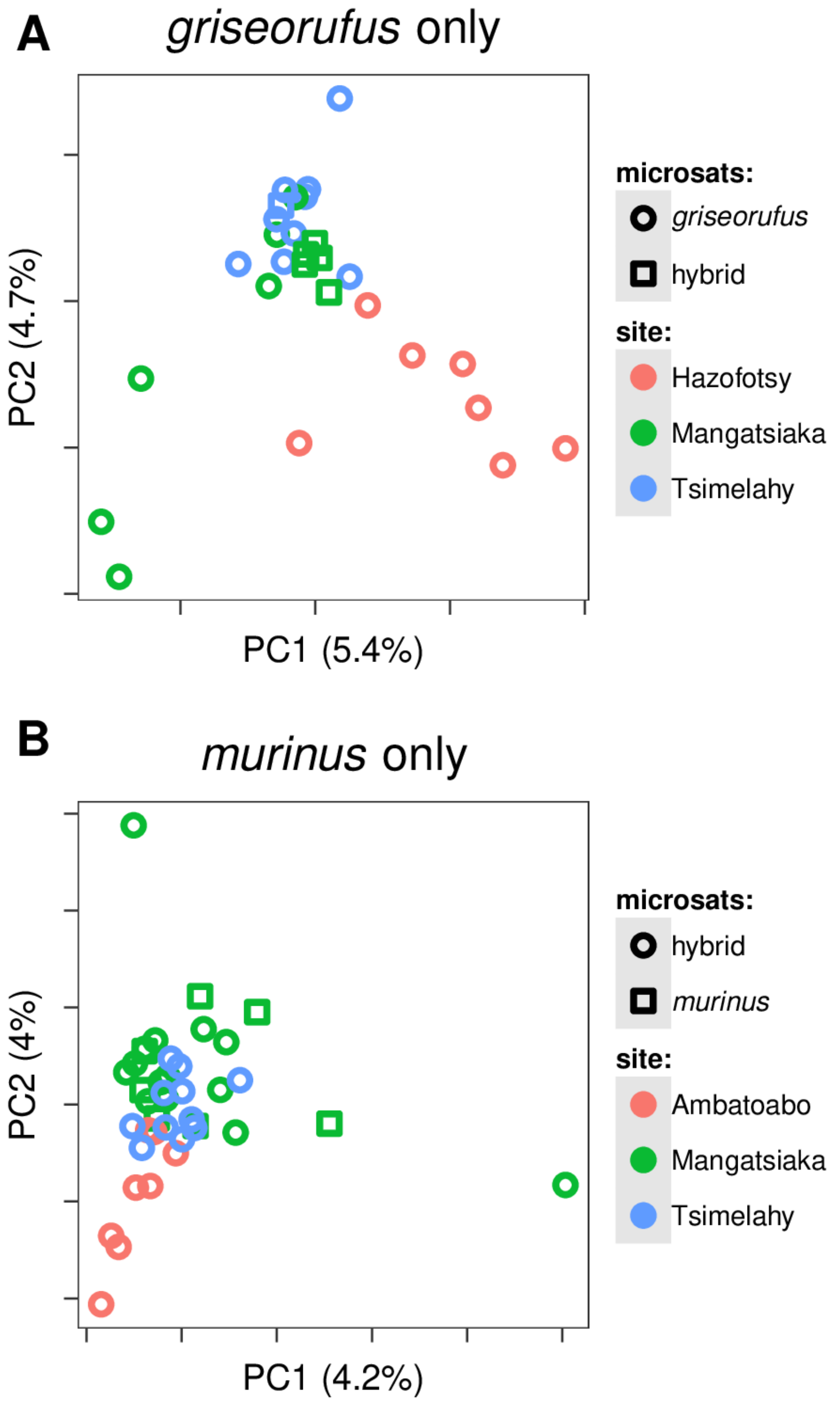

**Fig. S12: Within-species PCAs**
The “microsats” designation indicates how Hapke et al. (2011) classified the sample using microsatellites.

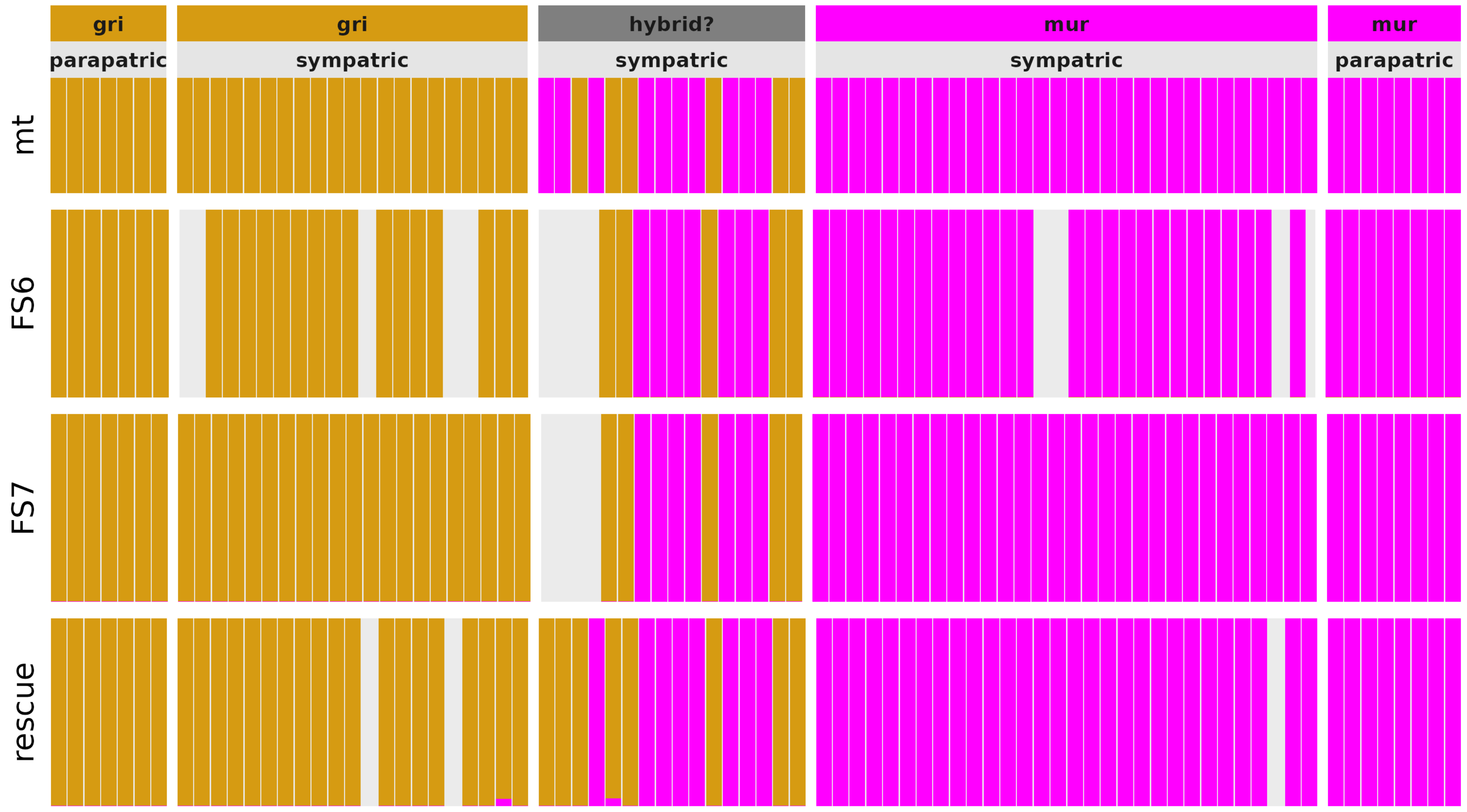

**Fig. S13: ADMIXTURE results for different filtering options**
Gray areas indicate that the samples in question were not present in the focal VCF file. Two samples (mhyb001 and mhyb002) only present in the “rescue” dataset show *griseorufus* nuclear DNA, but a *murinus* mitochondrial haplotype, in line with results from Lüdemann (2018). Two other samples (mhyb003 and mgri104) show <5% *murinus* ancestry in the “rescue” dataset, but these samples did not show any *murinus* ancestry in the FS6 and FS7 datasets, which contain many more SNPs
205 and are therefore more reliable.

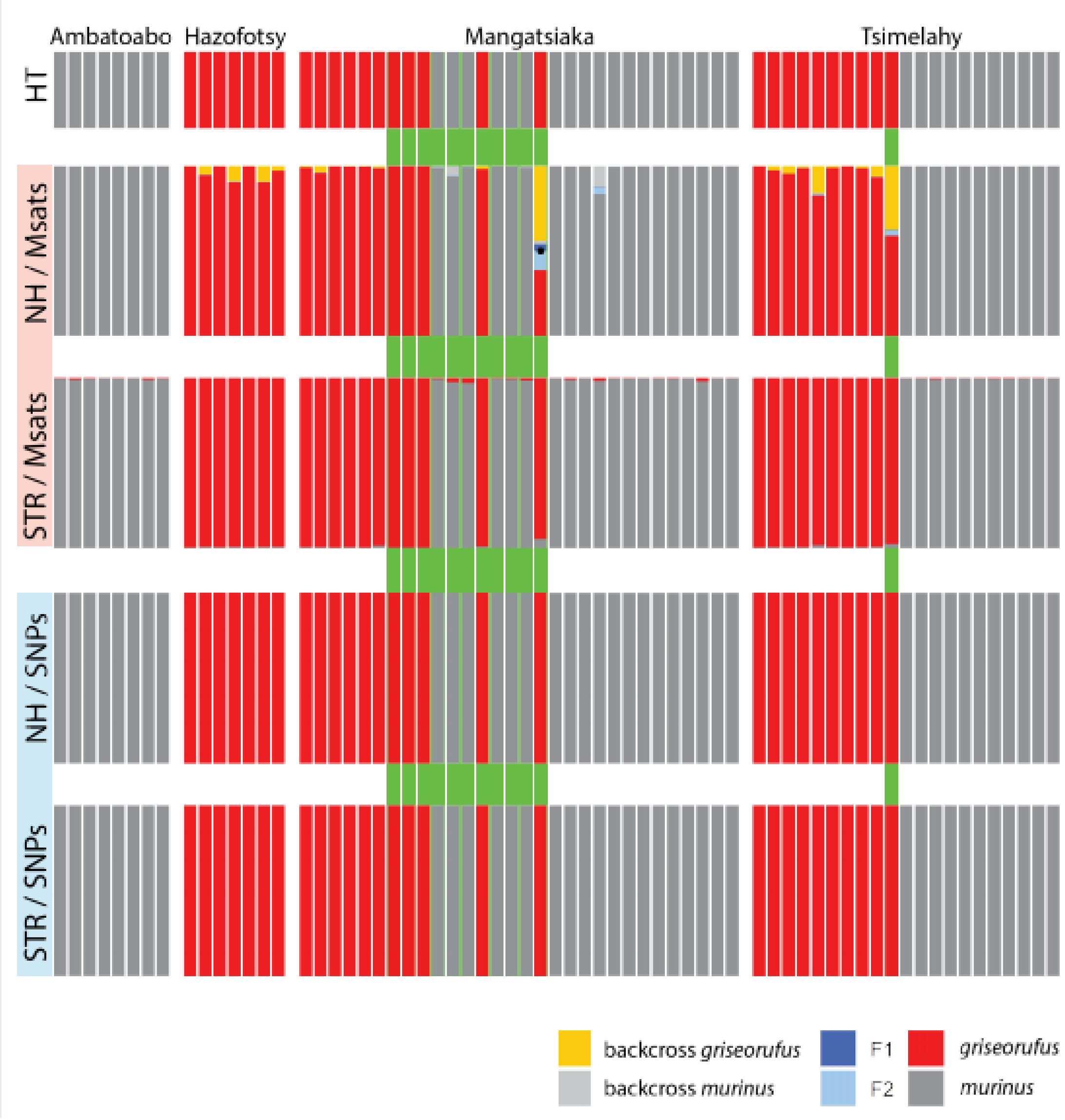

**Fig. S14: Re-analysis of microsatellite data for individuals with RADseq data.**
*Top row:* mitochondrial haplotype (HT).
*Next two rows:* microsatellites (Msats) analysed with NewHybrids (NH) and Structure (STR).
*Bottom two rows:* RADseq SNPs (SNPs) analysed with NewHybrids (NH) and Structure (STR).

While no hybrids are detected using RADseq data, NewHybrids identifies a single hybrid (black dot) using the microsatellite data, with several further *griseorufus* individuals showing non-significant signs of admixed ancestry (yellow ancestry).

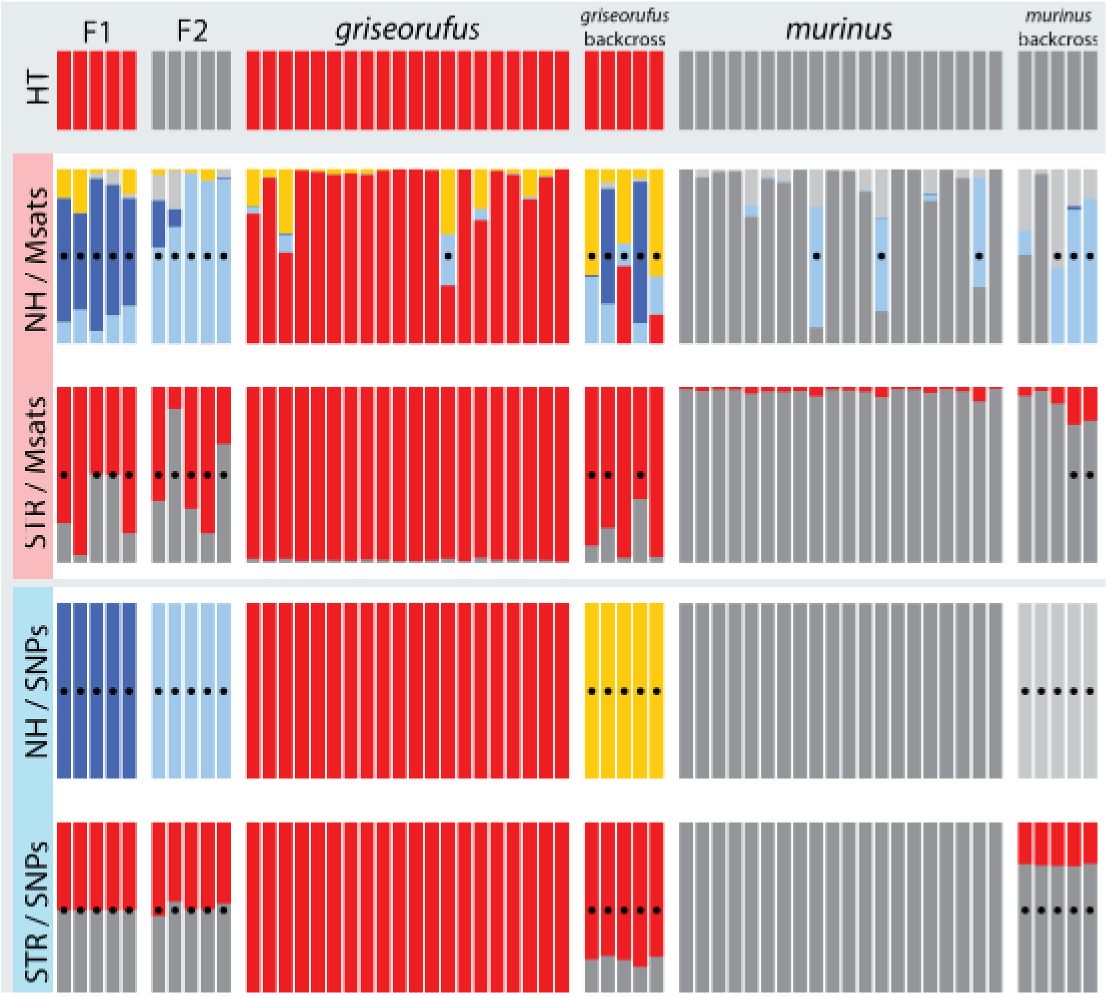

**Fig. S15: Assignment of simulated individuals using microsatellites and SNPs with Structure and Newhybrids.**
A black dot indicates that a given sample was classified as a hybris.
HT: mitochondrial haplotype, NH: NewHybrids, STR: Structure, SNPs: RADseq SNP, Msats: microsatellites.

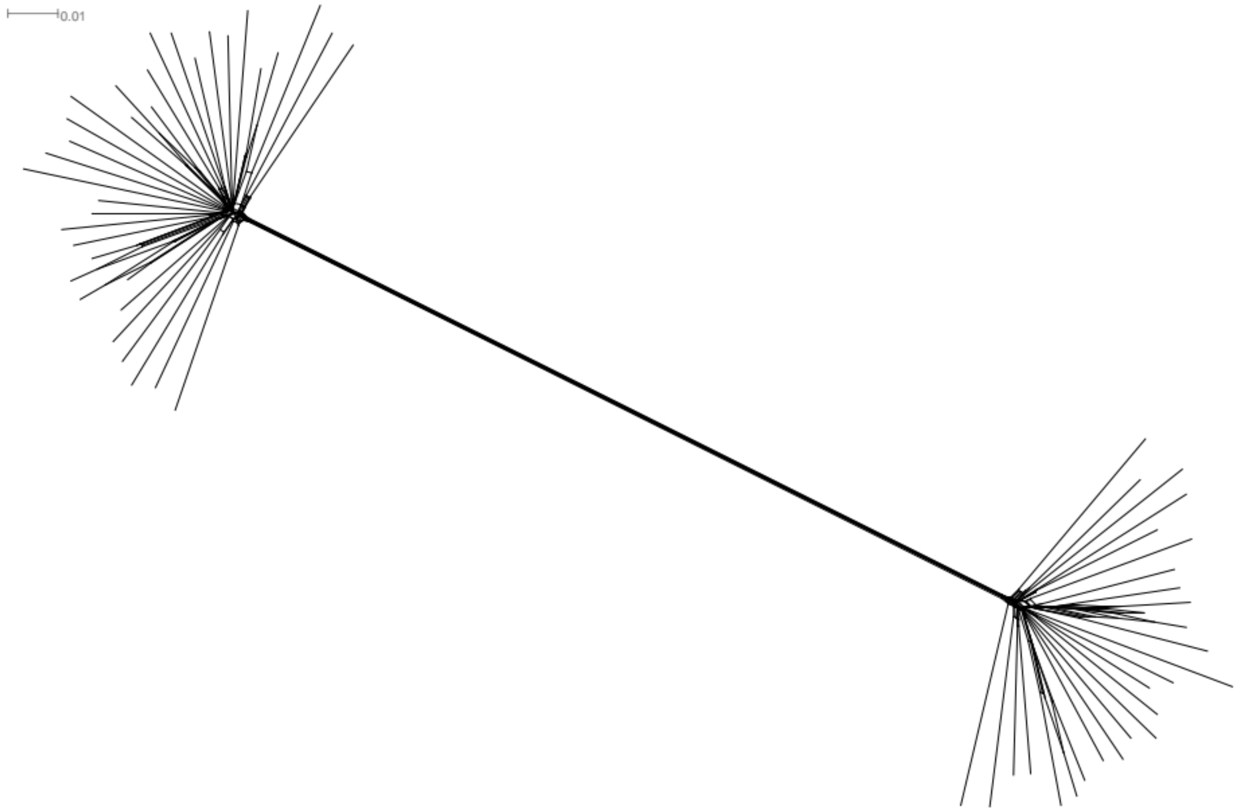

**Fig. S16: A SplitsTree NeighborNet phylogenetic network using only contact zone individuals.**

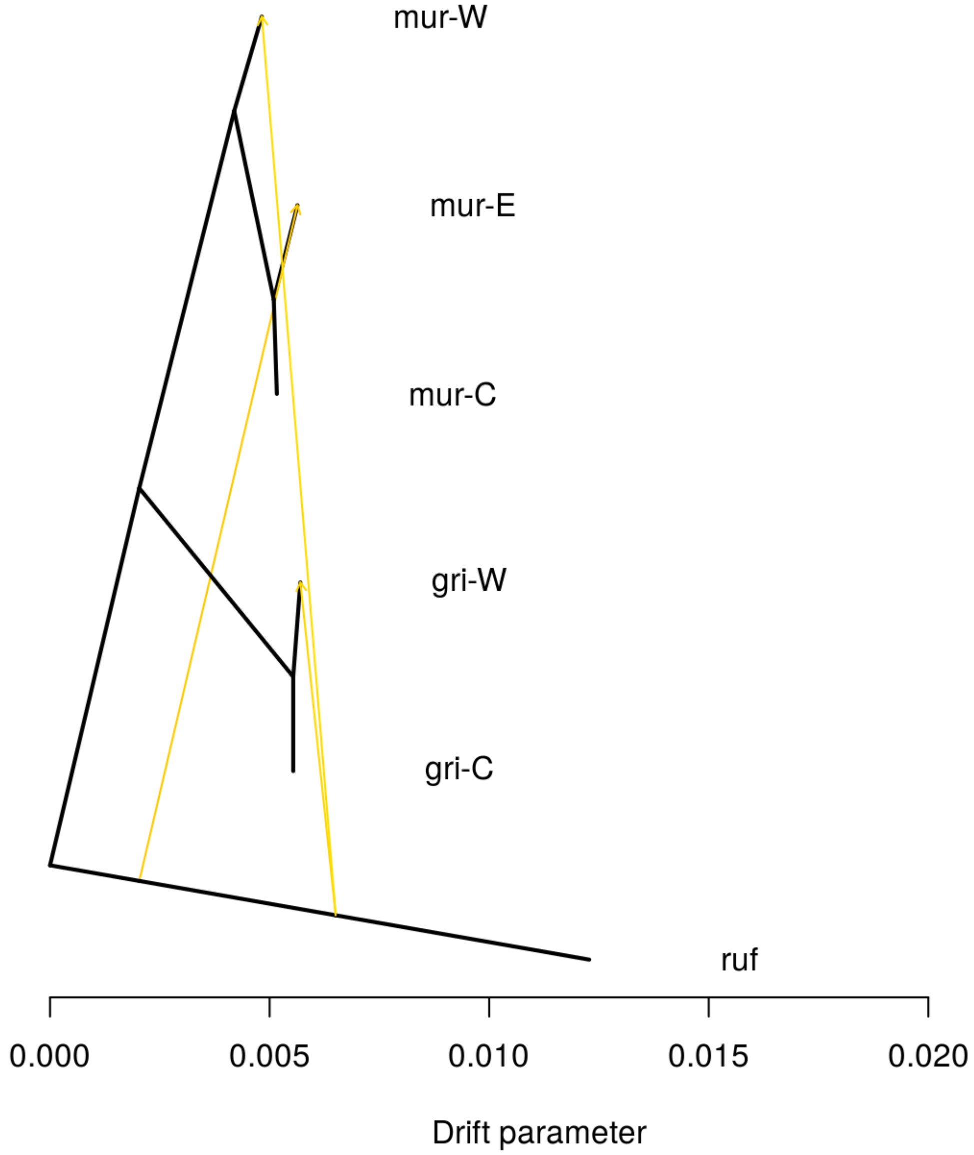

**Fig. S17: Significant migration edges using Treemix (with *M. rufus)*.**
All three migration edges are minor and involve *M. rufus*.

245

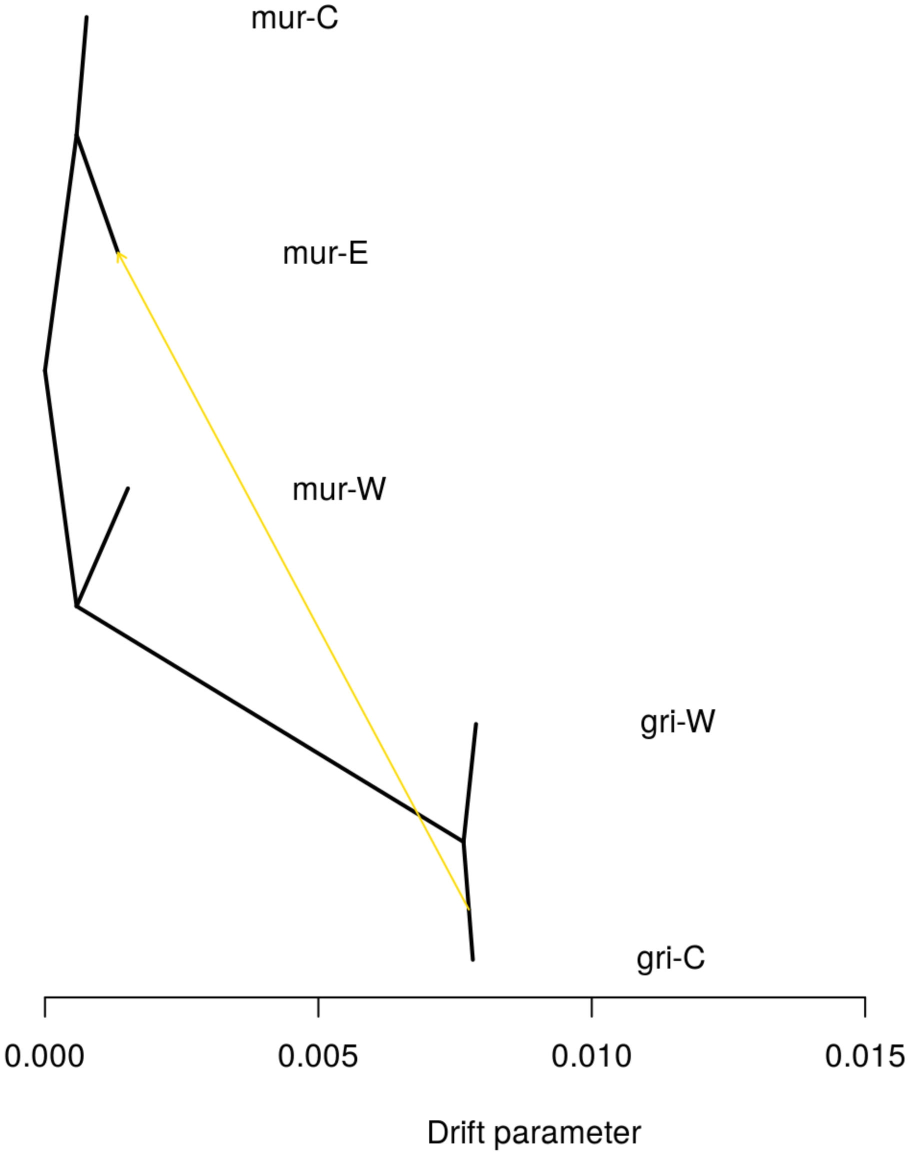

**Fig. S18: Significant migration edges using Treemix (without *M. rufus)*.**
